# Sparse mixed codes on shared manifolds for human-like spatial attention in artificial neural networks

**DOI:** 10.64898/2026.02.15.705979

**Authors:** Mingchen Liu, Fei Song, Bailu Si, Liqin Zhou, Ke Zhou

## Abstract

Spatial attention is often partitioned into endogenous, exogenous, and social forms, yet it remains unclear whether a single neural circuit can support all three and how their population codes are organized. Here we trained recurrent artificial neural networks (ANNs) with convolutional sensory front-ends on three classic cueing paradigms (central, peripheral, and gaze cues) to reproduce human reaction time (RT) profiles across cue–target onset asynchronies. Despite differences in sensory architecture and visual experience, all ANNs captured the full facilitation–inhibition time course for all three attention types. Model-based targeted dimensionality reduction (mTDR) revealed that cue- and choice-related activity in the advanced cognitive module evolved as rotations within a shared low-dimensional manifold, with angular deflections that mirrored the distinct temporal dynamics of endogenous, exogenous, and social attention. Attentional signals were encoded by highly sparse, distributed population activity: a small subset of recurrent units explained most task-related variance, was sufficient to recover human-like RT patterns after virtual lesioning, and became progressively sparser as training improved performance. At the same time, single unit responses displayed pervasive mixed selectivity, dominated by nonlinear conjunctions of cue type, cue direction, and validity, whose strength and heterogeneity robustly predicted model performance. Together, these results identified low-dimensional geometric rotations, sparse coding, and nonlinear mixed selectivity as core computational principles through which a single recurrent circuit could generate human-like temporal dynamics across endogenous, exogenous, and social orienting, and provided testable predictions for population-level mechanisms of spatial attention in the brain.

## Introduction

Selective attention prioritizes task-relevant or salient information while suppressing distractors, thereby gating sensory evidence to perception, working memory, and decision-making. Classical theories divide spatial selective attention into endogenous (goal-directed) and exogenous (stimulus-driven) forms, which differ in triggering conditions, temporal dynamics, and neural substrates (Corbetta & Shulman, 2002; Petersen & Posner, 2012; Posner, 1980; Vossel et al., 2014). Endogenous attention is voluntarily deployed by internal goals, emerges relatively slowly (approximately 300 ms after central cue onset) (Cheal & Lyon, 1991), and can be sustained without inhibition of return (IOR) (Muller & Rabbitt, 1989); anatomically, it engages the dorsal attention network (DAN), including the intreparietal sulcus (IPS) and frontal eye field (FEF)(Fox et al., 2006; Petersen & Posner, 2012). Exogenous attention, in contrast, is reflexively captured by salient, nonpredictive peripheral cues, with rapid, transient facilitation peaking at around 100 ms (Cheal & Lyon, 1991; Muller & Rabbitt, 1989), followed by IOR at cue-target onset asynchronies (CTOAs) longer than around 250 ms (Klein, 2000). This form of orienting is linked to a predominantly right-lateralized ventral attention network (VAN), encompassing temporoparietal junction (TPJ) and ventral frontal cortex (VFC)(Asplund et al., 2010; Fox et al., 2006; Shulman et al., 2010). Behaviorally, central and peripheral cues can exert independent, additive influences on information processing (Riggio & Kirsner, 1997).

Despite this canonical dichotomy, it remained unsettled whether endogenous and exogenous orienting constitute separable systems or partly overlapping computations within shared circuitry. Converging neuroimaging evidence pointed to both commonalities and dissociations. Event-related fMRI studies reported no reliable differences when directly contrasting endogenous and exogenous shifts, implicating a common frontoparietal network (Peelen et al., 2004). In parallel, Vossel et al. (2014) emphasized that the DAN and VAN were anatomically dissociable yet dynamically coupled, suggesting context-dependent interplay rather than strict independence. When goals and target locations were matched, both attention types robustly engaged the DAN, but regional contributions varied over time and across hemispheres depending on the orienting mode (Meyer et al., 2018). Consistently, single-unit decoding in macaque lateral PFC revealed partially shared neural resources supporting both voluntary and reflexive shifts, enabling a dynamic balance between internal goals and external demands (Di Bello et al., 2025). Causal evidence further demonstrated selective dissociations: lateral PFC lesion impaired endogenous but spare exogenous orienting (Rossi et al., 2007), whereas right parietal repetitive transcranial magnetic stimulation (rTMS) reduced exogenous orienting while leaving endogenous attention relatively intact (Hodsoll et al., 2009).

Social attention, exemplified by orienting to others’ gaze direction, also could not map cleanly onto endogenous or exogenous attention. Gaze cues were centrally presented yet trigger reflexive facilitation (Friesen & Kingstone, 1998), with benefits building more slowly (at about 200-300 ms CTOA), closer to endogenous timing (Driver et al., 1999). IOR was also markedly delayed, emerging at extended CTOAs (e.g., ∼1200 ms), much later than the IOR typically observed with peripheral cues (Frischen & Tipper, 2004; Jingling et al., 2015). Neuroimaging studies indicated that gaze and arrow cues engaged overlapping DAN/VAN circuitry (Joseph et al., 2015; Sato et al., 2009, 2016; Tipper et al., 2008; Uono et al., 2014), while social orienting additionally recruited occipito-temporal regions such as superior temporal sulcus/gyrus (STS/STG) involved in face and biological-motion processing (Gao et al., 2025; Hooker et al., 2003; Kingstone et al., 2004; Lockhofen et al., 2014; Wang et al., 2024). Other studies reported differential activation for social versus arrow cues within these networks (Engell et al., 2010; Hietanen et al., 2006), and patient with prefrontal damage exhibited impaired endogenous and gaze-based orienting but preserved peripheral orienting (Vecera & Rizzo, 2004, 2006). Split-brain findings further indicated that social orienting depended on lateralized cortical connections (Kingstone et al., 2000), whereas exogenous and endogenous attention had been linked to subcortical contributions (Gabay & Behrmann, 2014; Guedj & Vuilleumier, 2023). Together, these observations argue against a purely shared or fully distinct mechanism, and instead point to a domain-general frontoparietal control system coupled to partially specialized social representations.

Despite substantial progress, the computational relationship among endogenous, exogenous, and social attention remained poorly understood. Most empirical studies focused on one or two attention types in isolation and were constrained by the difficulty of measuring large-scale population dynamics for all three with high temporal and spatial resolution in the biological brain. As a result, it was still unclear whether different forms of spatial attention relied on distinct circuit modules, or instead arose from shared, high-dimensional population codes that were differentially configured across time and state space. Addressing this question required a framework that linked behavior, neural geometry, and population coding principles within a single model capable of performing all three attention tasks.

Artificial neural networks (ANNs) offered a complementary computational framework for exploring the mechanisms of attention. Beyond their established applications in modelling perception, memory, and decision-making (Masse et al., 2019; Richards et al., 2019; Roseboom et al., 2019; Sheahan et al., 2021; Ye et al., 2025), ANNs could be trained on multiple cognitive tasks concurrently regardless experimental time constraints, enabling systematic evaluation of the computational principles underlying multi-tasking (Dubreuil et al., 2022; Piwek et al., 2023; Yang et al., 2019). Here we developed several ANN models with biologically inspired visual front-ends and trained them simultaneously on endogenous, exogenous, and social cueing paradigms that reproduce human reaction time (RT) profiles across CTOAs. We then combined clustering, neural manifold analysis, and population-level representational characterization in the advanced cognitive module to address the following questions: (i) could a single recurrent architecture generate all three forms of spatial attention with human-like temporal dynamics; (ii) were endogenous, exogenous, and social orienting implemented by segregated functional clusters or by shared low-dimensional manifolds; and (iii) which population coding principles were most tightly linked to human-like attentional performance.

## Results

### 1. ANNs trained to exhibit human-like spatial attention behavior

We trained an ANN model on three classical spatial precueing paradigms □ central, peripheral, and social (Fig. 1a) □ designed to capture endogenous, exogenous, and social attention, respectively. Cue validity (CV) was set to 70% for central cue, and 50% for peripheral and social cues. The experimental timing parameters were matched to established human paradigms (see *Methods*). The design aimed to establish an ANN model capable of reproducing typical human attention dynamics across CTOAs: slow but sustained facilitation for endogenous attention, fast facilitation followed by IOR for exogenous attention, and slow facilitation with a later-onset IOR for social attention (Fig. 2a).

**Fig. 1.**
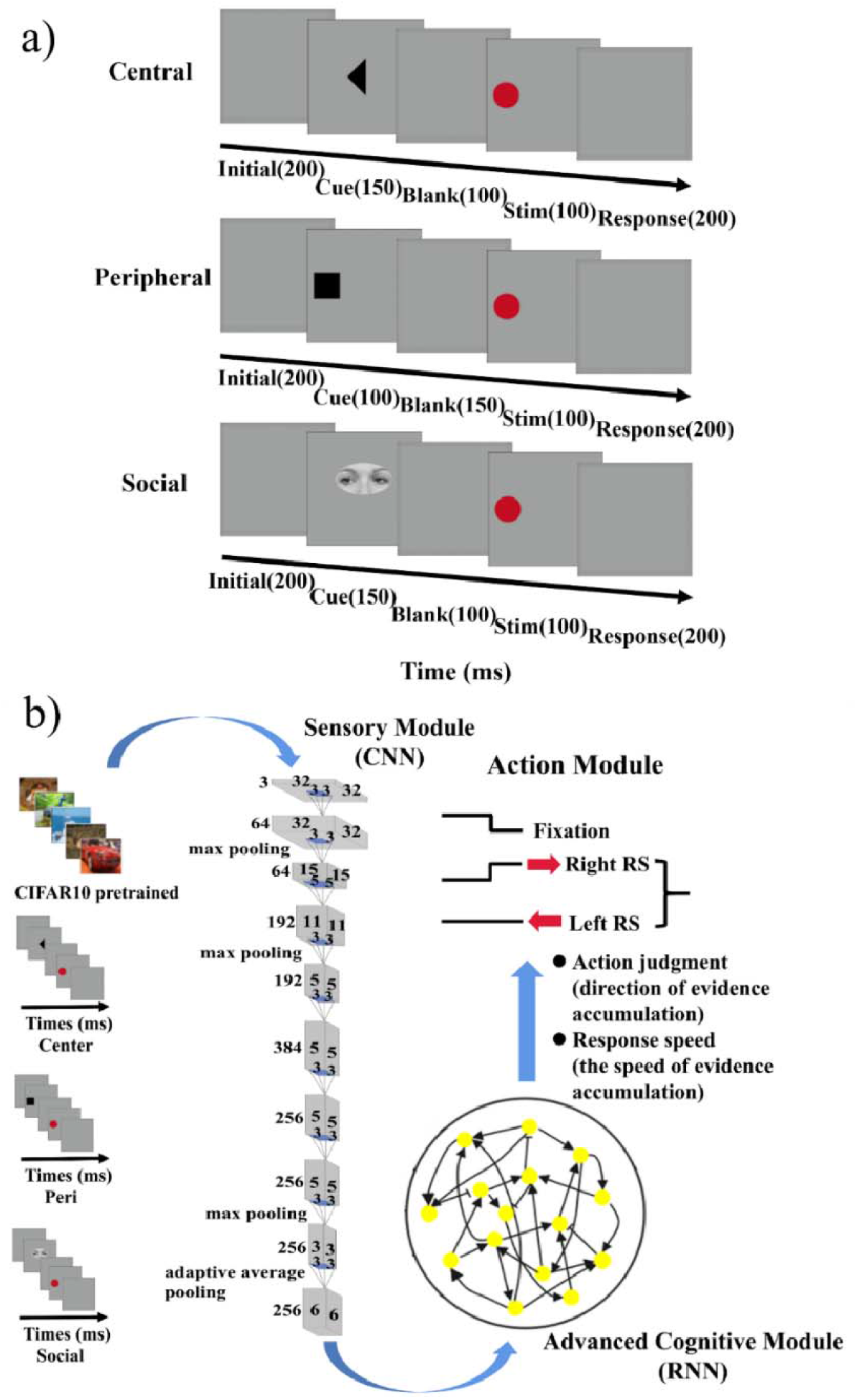
Task design and architecture of the Pretrained AlexNet model. (a) Example trial sequences for the central, peripheral, and social cueing paradigms at CTOA = 250 ms. Each trial consisted of an initial blank period, presentation of a spatial cue (central arrow, peripheral square, or gaze stimulus), a blank interval, target onset, and a response window. (b) The network architecture of the Pretrained AlexNet model. The model comprised a sensory module, an advanced cognitive module, and an action module. The sensory module was an AlexNet convolutional network pretrained on CIFAR-10 and kept fixed during attention training. Trial images were streamed through the sensory module into the advanced cognitive module. The action module included three units: a fixation unit and two units encoding reaction speed (RS) for right- and left-side choices.

**Fig. 2.**
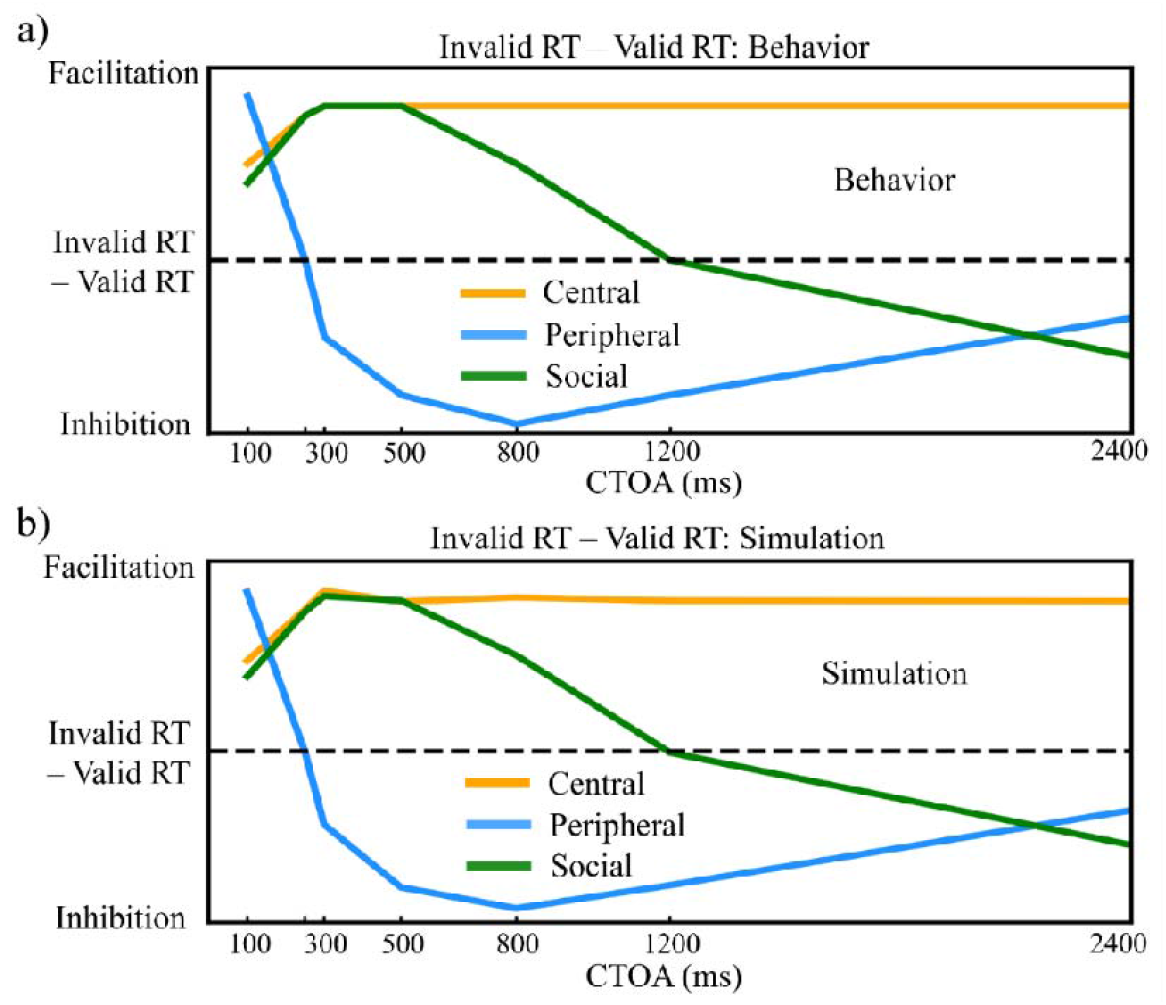
Human and model spatial attention effects. (a) Empirical attention effects for central, peripheral, and social cueing tasks. The ordinate showed invalid RT minus valid RT; positive values indicated facilitation, whereas negative values indicated inhibition. (b) Simulated attention effects from the Pretrained AlexNet model plotted in the same format.

The ANN model consisted of three modules: a sensory module, a cognitive module, and an action module (Fig. 1b). The sensory module was an AlexNet (Krizhevsky et al., 2017) pretrained on CIFAR10, providing natural visual experience analogous to the human visual ventral stream. The cognitive module contained 256 leaky recurrent units, using for higher-level cognitive processing. This configuration was referred to as the Pretrained AlexNet model. The action module was to determine whether response was made and the extent of the response.

Training results showed reliable convergence, with loss stabilizing after ∼400 epochs (Fig. S1). The Pretrained AlexNet model not only achieved perfect accuracy in RT prediction but also closely reproduced both the magnitude and temporal profile of human attention effects (i.e., the RT differences between valid and invalid conditions) (Fig. 2b, Fig. S2). To assess training robustness, we trained 50 independent instances of the Pretrained AlexNet model with identical hyperparameters, initialization, and optimization settings. All instances showed highly similar loss trajectories (Fig. S3a) and replicated the spatial-attention effects with minimal across-instance variability (Fig. S3b), confirming stable learning dynamics.

To further explore the general neural representational characteristics of spatial attention and disentangle the contributions of pretraining experience and network architecture on simulating performance, we further implemented other five ANNs with distinct visual experience and sensory module architectures: (1) a Pretrained ResNet model (Fig. 3b) to test the influence of alternative sensory architecture; (2) a Stimulus AlexNet model (Fig. 3c), pretrained on experimental stimuli with frozen sensory weights during attention task training, to evaluate task-specific visual experience; (3) an End-to-end AlexNet model (Fig. 3d), jointly trained the modules without pretraining, to assess the effect of concurrent sensory-cognitive learning; and two randomly initialized models with fixed weights, (4) Random AlexNet model (Fig. 3e) and (5) Random ResNet model (Fig. 3f), to control for the absence of visual experience.

**Fig. 3.**
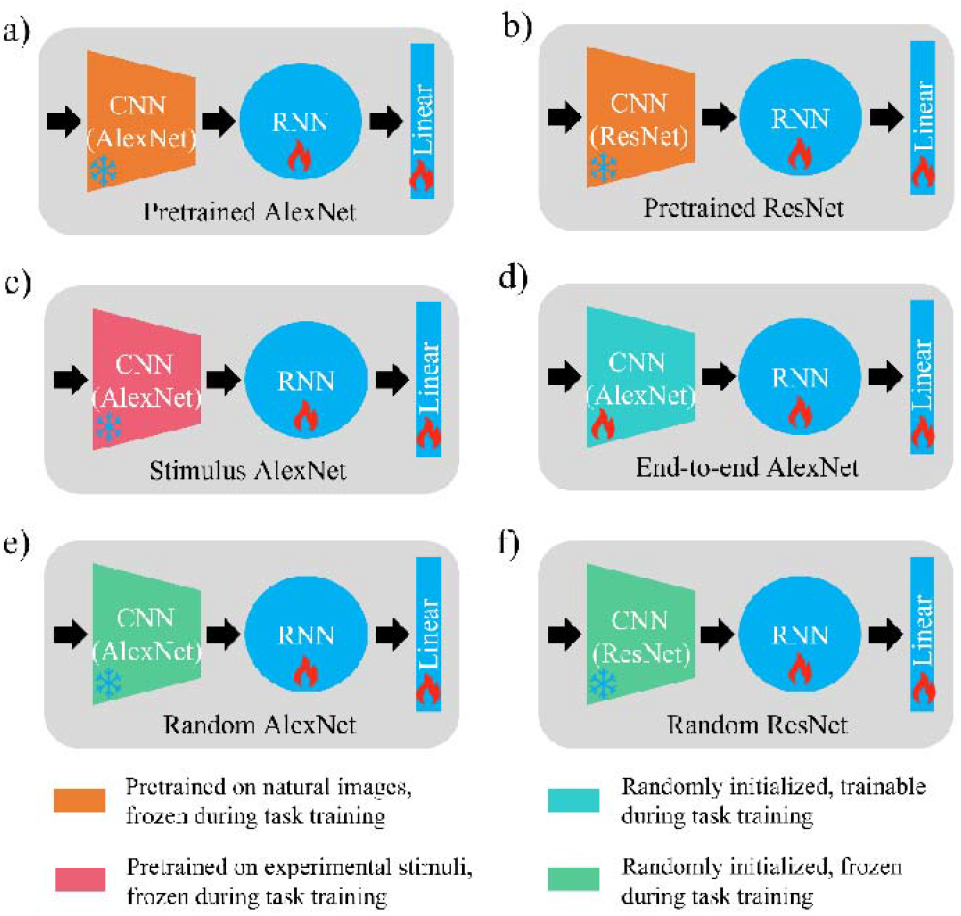
Architectures and training regimes of comparison models. (a) Pretrained AlexNet; (b) Pretrained ResNet; (c) Stimulus AlexNet; (d) End-to-end AlexNet; (e) Random AlexNet; (f) Random ResNet.

All five ANNs learned to quantitatively reproduce human spatial attention effects (Fig. S4), though they required different number of epochs to reach convergence (Fig. S1). These findings established that, despite architectural and experiential differences, the ANNs could acquire human-like spatial attention dynamics, providing a foundation for investigating how cognitive modules develop shared and task specific neural representations of distinct types of spatial attention.

### 2. Neural geometry in advanced cognitive module

To investigate how cue- and choice-related signals are represented and interact within the advanced cognitive module, we used model-based targeted dimensionality reduction (mTDR) method to project high-dimensional population activity under each cue type and CTOA onto orthogonal low-dimensional subspaces defined by “cue” and “choice” axes.

At short CTOAs (100-300 ms), trajectories remained nearly symmetric around the choice axis, indicating minimal directional bias when cue-target delays were brief (Fig. 4a-c). As the interval increased (500-800 ms CTOAs; Fig. 4d-e), trajectories diverged systematically across cue types. For central and social cues, valid trials gradually shifted toward the cued location, while invalid trials deflected oppositely; for peripheral cues, this pattern was reversed. At longer CTOAs (1200-2400 ms; Fig. 4f-g), central and peripheral cues preserved their earlier patterns, whereas the social-cue trajectory gradually reversed direction, reflecting a late inhibitory phase. This transition from early symmetry to late cue-dependent divergence suggests a gradual integration of spatial context into the internal state representation of the cognitive module, consistent with the increasing variance explained by the cue axis across CTOAs (Table S1).

**Fig. 4.**
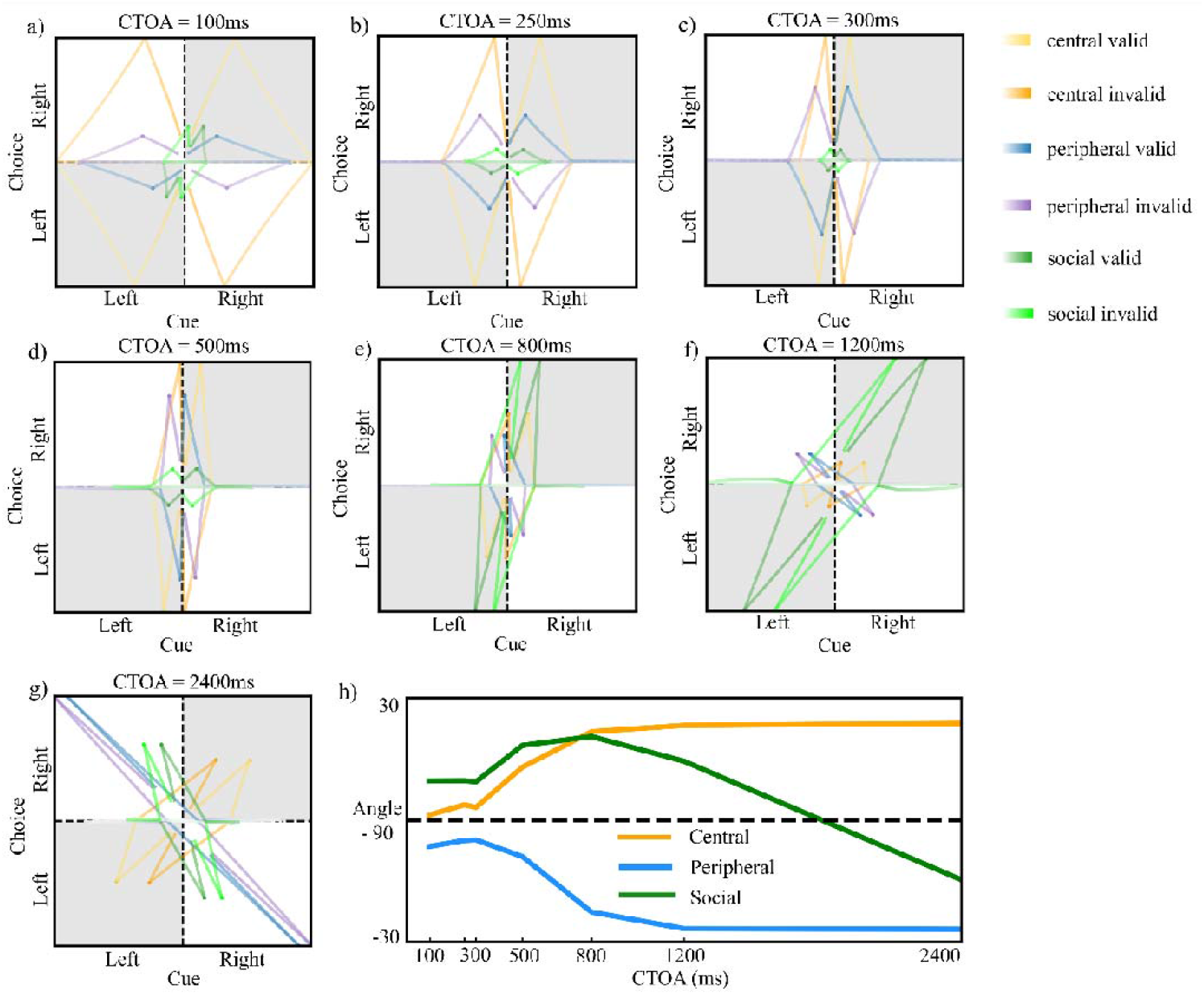
Neural geometry in the advanced cognitive module of the Pretrained AlexNet model. (a–g) Trial-averaged population activity projected onto cue (horizontal) and choice (vertical) axes for central (yellow), peripheral (blue/purple), and social (green) cues at different cue–target onset asynchronies (CTOA = 100, 250, 300, 500, 800, 1200, 2400 ms). Gray shading in the first and third quadrants marks states in which choice activity was aligned with the cued direction (facilitation). (h) Deflection angle between valid and invalid trajectories relative to the neutral axis (90°) as a function of CTOA.

To quantify these neural biases, we computed the acute angle between the symmetry axis of cue-specific trajectory and the vertical choice axis (Fig. 5b). The resulting angular deflections closely paralleled the behavioral facilitation–inhibition profile (Fig. 2a and Fig. 4h). Central and social cues exhibited progressively larger positive deviations that peaked at 500–800 ms, followed by a reversal toward negative angles for the social cue at longer CTOAs, reflecting delayed inhibition. In contrast, peripheral cues showed early negative deflections that intensified and stabilized, indicating rapid inhibition. Although these neural trajectories qualitatively mirrored behavioral trends, they did not fully capture the complete facilitation–inhibition time course, likely due to shifting contributions of cue-versus choice-related variance over time (Table S1).

**Fig. 5.**
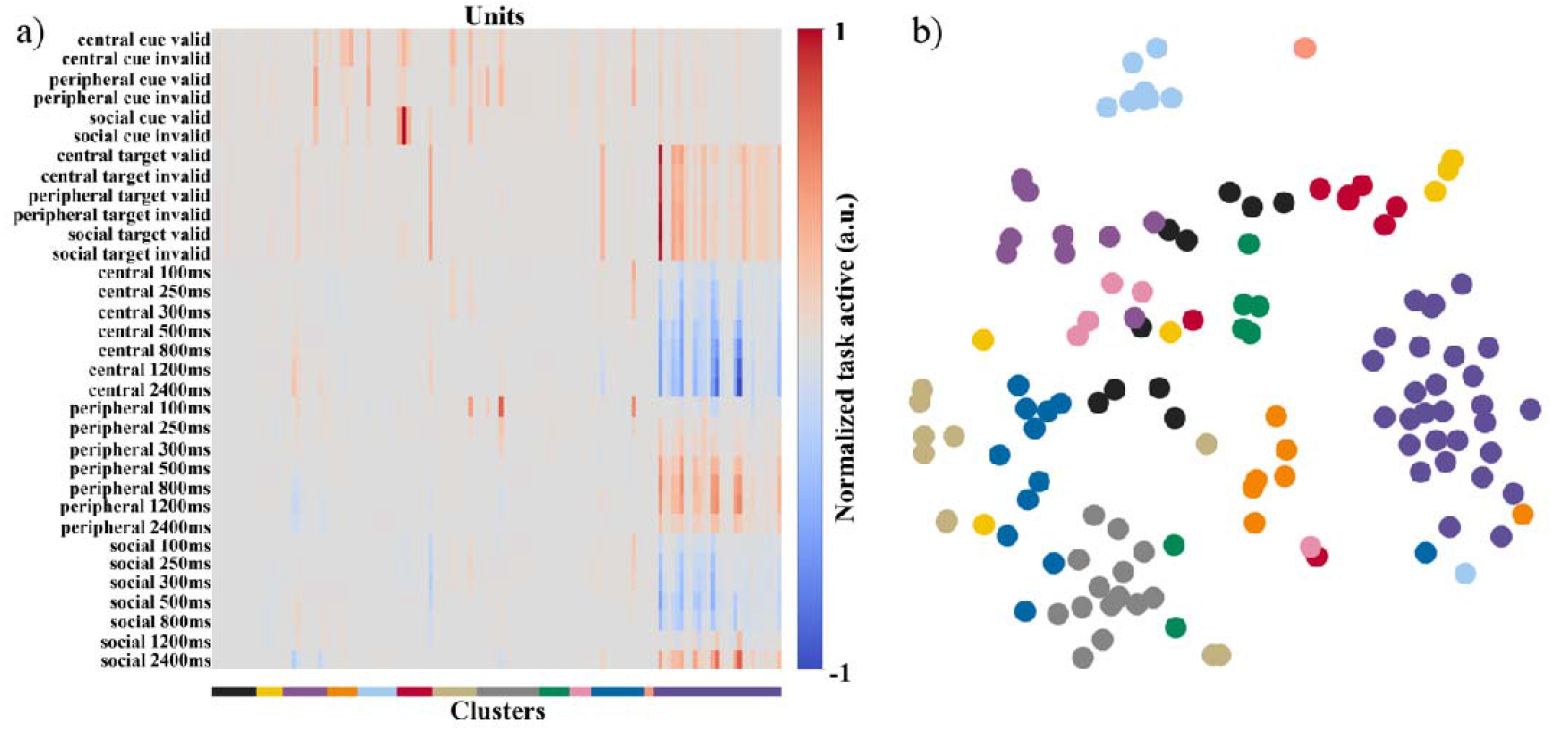
Functional clustering in the advanced cognitive module of the Pretrained AlexNet model. (a) Heat map of normalized activation for each recurrent unit, sorted by k-means cluster (clusters 1–13 on the horizontal axis) across 33 task features. (b) t-SNE embedding of the same units, with points colored by cluster identity, illustrating the weak separation among putative functional groups.

We next examined how pretraining experience and network architecture influenced these state-space dynamics (Fig. S5c-g). Three consistent patterns emerged. First, pretrained networks (Pretrained AlexNet and ResNet) exhibited robust, biphasic facilitation-inhibition trajectories that closely resembled human RT curves. Second, Stimulus-trained and End-to-end models reproduced central and peripheral attention dynamics but failed to capture the late inhibitory reversal observed for social cue, highlighting the critical role of natural visual experience. Third, randomly initialized networks exhibited attenuated or distorted deflection patterns, yet deeper architecture (e.g., Random ResNet) partially recovered the temporal organization of pretrained models.

Together, these findings demonstrated that both large-scale natural image pretraining and network architecture jointly shaped the population geometry underlying spatial attention by organizing cue–choice signals within shared low-dimensional manifolds, enabling a single recurrent circuit to generate human-like temporal dynamics of endogenous, exogenous, and social spatial attention after limited task-specific training.

### 3. No specific functional clusters encode spatial attention tasks

To examine whether the cognitive module exhibited functionally organized subpopulations, we applied k-means clustering on unit activation patterns. For each unit, three activation features across cue types, validity conditions, and task periods were extracted: (1) mean activation during the cue period, (2) mean activation during the target and response periods, and (3) the difference in mean activation between valid and invalid cues at each CTOA.

In the Pretrained AlexNet model, clustering analysis revealed no evidence of well-separated functional groups. Silhouette scores remained consistently below 0.3 across all tested cluster numbers (2-20; Fig. S6a), indicating weak cluster structure and highly overlapping selectivity among units. Although certain clusters (e.g., cluster 4-9) showed enhanced response during the cue period and another (cluster 11) during the target and response periods, these subgroups exhibited broad and partially overlapping response profiles rather than distinct condition-specific representations (Fig. 5). Most clusters contained mixed units responsive to multiple cue types, and only a small subset of neurons exhibited uniformly high activation across all conditions, reflecting a sparse yet distributed representation of attention-related signals.

Across all other network variants, a similar pattern emerged: Silhouette scores remained below 0.3 (Fig. S6b–f), and their activity heatmaps showed highly interleaved activation patterns across conditions and periods (Fig. S7). These findings confirmed that units in all models operated as partially independent components within a shared representational space rather than forming coherent, modular clusters.

Collectively, these results indicated that the cognitive module developed only minimal functional specialization. Instead, spatial attention was encoded through distributed, sparse, mixed-selective population codes—a hallmark of flexible neural computation observed in biological cortical circuits. Such partially overlapping representations might allow the cognitive module to flexibly integrate information across cue types and time scales, supporting adaptive attentional processing.

### 4. Representation sparsity in advanced cognitive module

To assess how efficiently spatial attention is represented in the advanced cognitive module, we quantified the sparsity of population activity and examined its relationship to behavioral performance.

In the Pretrained AlexNet model, population activity was highly sparse and distributed rather than clustered into distinct functional groups. A small subset of units captured most of the task variance: 34 units (13.28% of all units) accounted for 95% and 57 units (22.27%) for 99% of the variance across cue type and validity conditions (Fig. 6a). Virtual ablation confirmed the functional sufficiency of this subset—when restricted to these units, the model maintained near-identical loss values (Fig. 6b) and reproduced the complete facilitation–inhibition behavioral profile (Fig. 6c-d). Consistent with this, as units were incrementally added in order of explained variance, the three spatial-attention effects emerged and sharpened in parallel, and the complete facilitation–inhibition pattern was recovered only after including the subset that accounted for most of the variance (Fig. S8). These results indicated that an overlapping, compact set of high-level units could support complex attentional computations with high fidelity and efficiency.

**Fig. 6.**
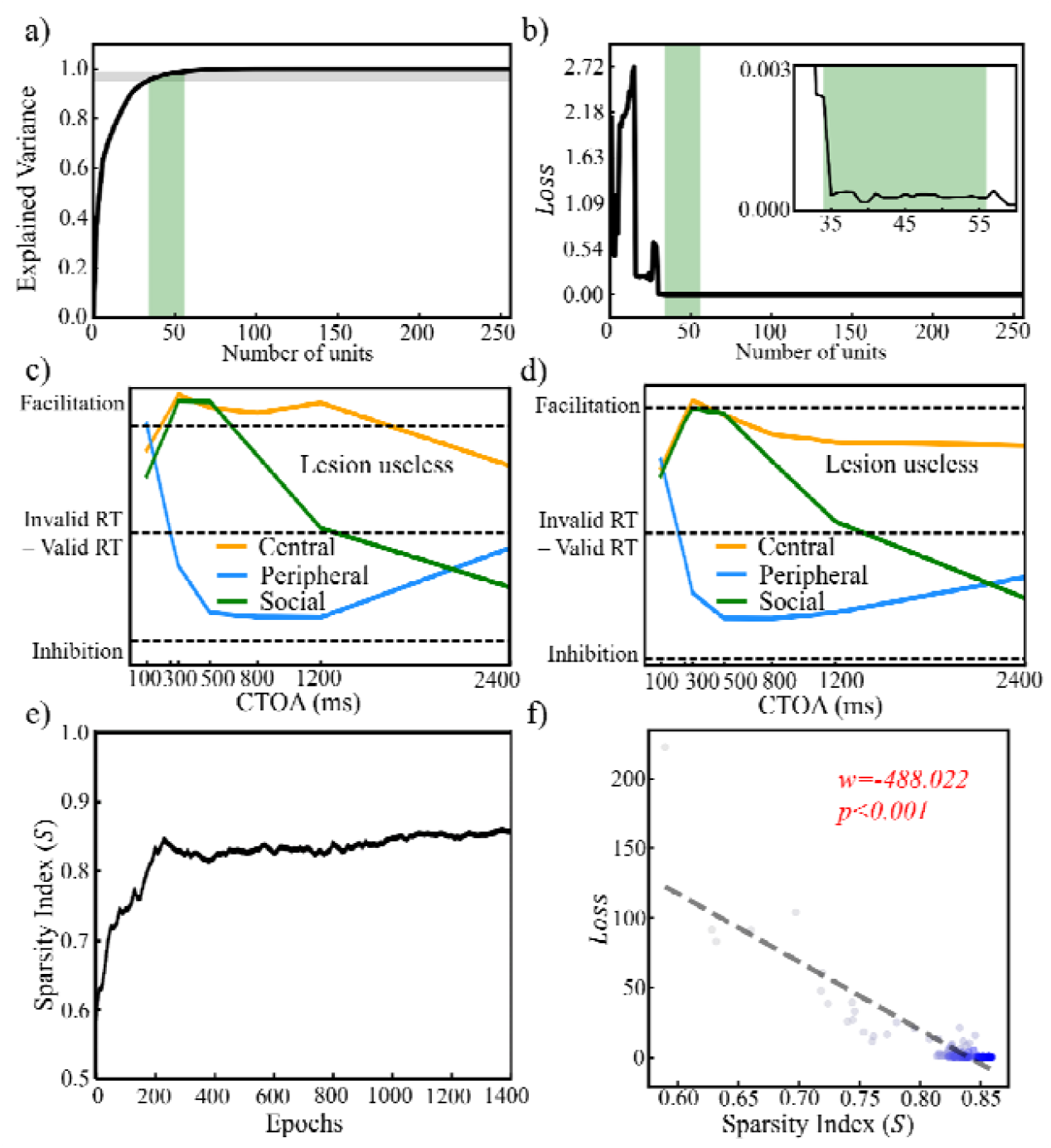
Sparse population code in the advanced cognitive module of the Pretrained AlexNet model. (a) Cumulative variance explained as a function of the number of recurrent units included. The black curve shows the model, horizontal gray lines mark 95% and 99% of total variance, and the green band indicates the range of units (34–57) required to reach these levels. (b) Model performance () when only the top k variance-contributing units were retained. The inset zoomed the range of 30–100 units; the green band again marked the 34–57 unit range. (c,d) Simulated attention effects after restricting the network to units explaining (c) 95% or (d) 99% of the variance. The facilitation–inhibition profiles closely matched those of the intact model, indicating that lesioning the remaining units has little impact. (e) Evolution of the sparsity index of the advanced cognitive module across training epochs. (f) Relationship between model performance () and the sparsity index. Each point corresponded to one training checkpoint (color denoted training progress, with darker points indicating later epochs); the dashed line showed the best-fitting linear regression.

The sparsity index of the Pretrained AlexNet cognitive module was 0.891±0.003, comparable to values observed in primate cortex, including the monkey inferior temporal cortex (0.75-0.78; (Kreiman et al., 2006)), temporal lobe (0.78; (Perrodin et al., 2011)), and orbitofrontal cortex (0.73-0.77; (Kadohisa et al., 2004)). This congruence suggested that the network, analogous to biological systems, accomplish spatial attention through a compact, energy-efficient code. During training, both behavioral fit (Fig. S9) and sparsity (Fig. 6e) increased steadily before saturating. Across training epochs, sparsity strongly and negatively predicted model loss (w = –488.022, p < 0.001; Fig. 6f), explaining about 90% of its variance. Thus, emergent sparsity constrained population activity onto a low-dimensional manifold, promoting efficient information encoding and precise behavioral output.

Across models, sparsity levels were uniformly high but systematically modulated by visual experience: Stimulus AlexNet (0.949±0.002) > Pretrained ResNet (0.896± 0.001) > Pretrained AlexNet (0.891±0.003) > End-to-end AlexNet (0.863±0.005) > Random ResNet (0.837±0.005) > Random AlexNet (0.803±0.005). Models with visual experience required substantially fewer active units to explain comparable variance (e.g., 20–30 units for the Stimulus AlexNet versus 58–102 units for random models to capture 95-99% of the variance; Fig. S10). Virtual ablation confirmed this efficiency (Fig. S10): pretrained and task-trained models maintained performance with small, compact unit subsets, whereas untrained networks required larger and more redundant populations.

Longitudinal analysis further revealed that task-driven optimization consistently increased sparsity across architectures, stabilizing after several hundred epochs (Fig. S11). Across all trained models and the Random ResNet, sparsity again negatively predicted model loss (w < −1110.838, p < 0.001; Fig. S12), except for the Random AlexNet (w = −74.299, p = 0.103). This negative association, consistent with findings in the Pretrained AlexNet, suggested that increased sparsity reliably facilitates performance improvement and the emergence of human-like spatial attention behavior.

Together, these findings identified sparse coding as one of the key organizational principles for flexible spatial attention. Visual experience enhanced both the degree and efficiency of sparsity, enabling the ANNs to achieve human-like attentional performance through compact, energy-efficient population representations.

### 5. Mixed selectivity in advanced cognitive module

Flexible spatial attention requires neural populations capable of encoding multiple task variables within a limited number of units. We therefore asked whether the advanced cognitive module developed high-dimensional, mixed-selective codes for cue type, cue direction, and validity. For each unit, we quantified its selectivity along these three factors and decomposed the response into linear (additive) and nonlinear (interactive) mixed components; we also tracked a population-level mixed-selectivity preference *I*_*mixed*_, which reflected the balance between linear and nonlinear tuning.

Across all six models, mixed selectivity emerged robustly and was dominated by the nonlinear component (Fig. 7). In the Pretrained AlexNet, Pretrained ResNet, and Random ResNet, the mean variance explained by nonlinear mixed selectivity (*Mean*_*nlin*_) rose sharply from near zero during early training and then saturated at a high level, whereas in the Stimulus AlexNet, End-to-end AlexNet, and Random AlexNet, *Mean*_*nlin*_ increased more gradually. This pattern indicated that both large-scale natural image pretraining and network architecture accelerated the formation of nonlinear mixed codes, but were not strictly required for them to arise. By contrast, the mean variance explained by the linear component (*Mean*_*lin*_) remained small and only slowly increased with epochs in all models. For both components, the across-unit variability—captured by the widths of the 95% confidence intervals ( *ΔCI*_*lin*_ and *ΔCI*_*nlin*_) — expanded over training, reflecting growing diversity in how individual units combine task variables. Aside from a brief transition early in training, *I*_*mixed*_ remained positive and close to 0.5 throughout learning, indicating a stable bias toward nonlinear coding without a major shift in the overall balance between linear and nonlinear contributions (Fig. S13).

**Fig. 7.**
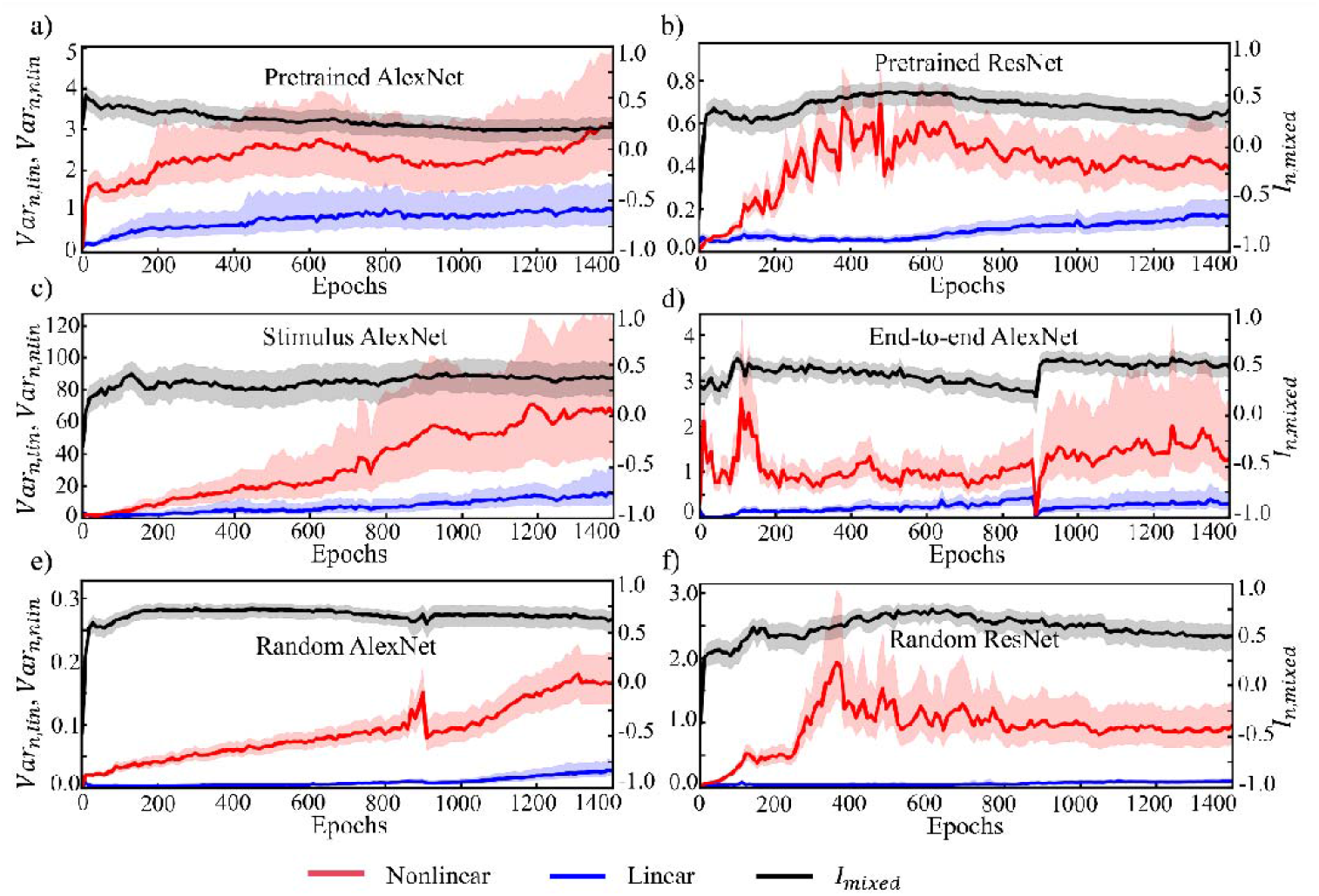
Mixed selectivity in the advanced cognitive module across training. (a) Pretrained AlexNet; (b) Pretrained ResNet; (c) Stimulus AlexNet; (d) End-to-end AlexNet; (e) Random AlexNet; (f) Random ResNet. For each model, the solid red curve showed the mean variance explained by the nonlinear mixed-selectivity component, and the solid blue curve showed the mean variance explained by the linear component. The solid black curve denoted the population-level mixed-selectivity preference,. Shaded regions indicated 95% CIs across units.

We next associated these linear and nonlinear mixed selectivity related measures to behavioral performance (Fig. 8). In every model, all four metrics (,,,), were negatively correlated with model loss (all slopes < 0, ≤ 0.001, except for in the End-to-end AlexNet). Thus, increases in either the strength or heterogeneity of mixed selectivity reliably accompanied reductions in model loss. Although linear metrics spanned a smaller numerical range and therefore exhibited steeper regression slopes, nonlinear components carried substantially more variance overall and were strong predictors of model loss.

**Fig. 8.**
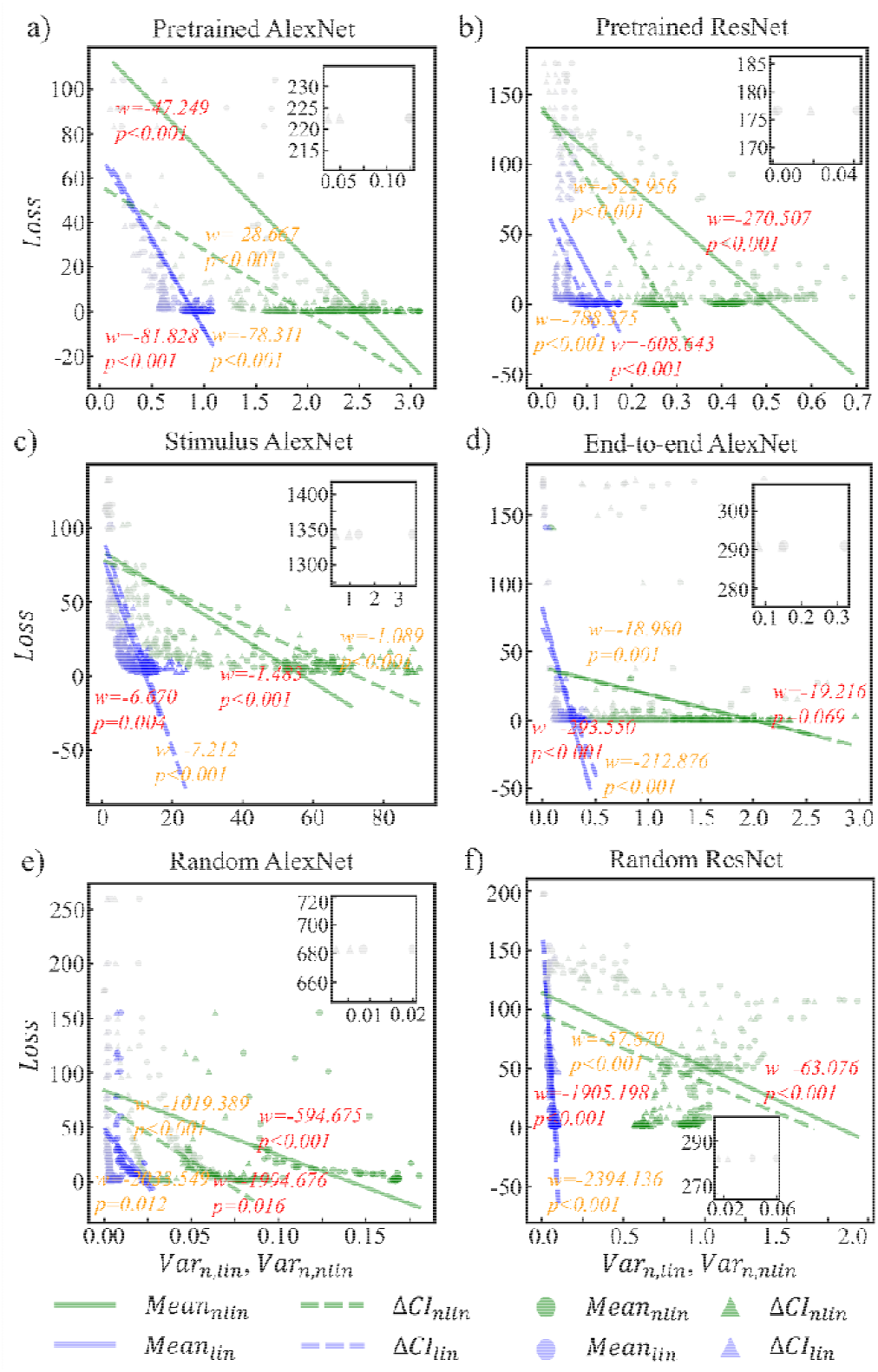
Relationship between model performance and mixed selectivity in the advanced cognitive module. (a) Pretrained AlexNet; (b) Pretrained ResNet; (c) Stimulus AlexNet; (d) End-to-end AlexNet; (e) Random AlexNet; (f) Random ResNet. Each point corresponded to one training checkpoint (every 10 epochs), with darker colors indicating later epochs. Green symbols and lines denoted the nonlinear component, and blue symbols and lines denoted the linear component. Circles indicated the population mean variance explained (,); triangles indicated the width of the 95% CI (,). Red text denoted the component, and orange text denoted the component.

Taken together, these results indicated that high-dimensional mixed selectivity, particularly its nonlinear component, served as a key organizing principle of the advanced cognitive module. Training sculpted a relatively small subset of units to encode conjunctions of cue type, cue direction, and validity. The subsequent increase in both the magnitude and diversity of these conjunctive codes closely tracked the emergence of human-like spatial-attention behavior.

## Discussion

Here we trained recurrent ANNs with biologically inspired visual front-ends to capture the behavioral signatures and neural population dynamics of endogenous, exogenous, and social spatial attention. Across the three cueing paradigms, the ANNs, including Pretrained AlexNet model, reproduced the full facilitation–inhibition profile of human RTs and, at the same time, developed strongly overlapping population codes with four key characteristics: cue- and choice-dependent rotations in a shared low-dimensional state space, a lack of sharply segregated functional clusters, highly sparse population activity, and pronounced mixed selectivity. Together, these features provided a unified geometric description of spatial attention in recurrent networks and illustrated how rich sensory priors, combined with architectural constraints, shaped human-like attentional computations.

Our manifold analyses showed that endogenous, exogenous, and social attention were implemented by cue- and choice-dependent rotations within a shared low-dimensional subspace. As CTOA increased, neural trajectories in the advanced cognitive module gradually rotated along cue and choice axes, transforming initially symmetric post-target states into strongly biased, cue-specific geometries that closely paralleled the facilitation–inhibition profile in behavior. This pattern was consistent with recent work using mTDR in frontal cortex, where stimulus, choice, and context were encoded in partially orthogonal subspaces whose relative contributions evolved over time rather than being statically multiplexed (Aoi et al., 2020; Pagan et al., 2025). Manifold analyses in both biological and artificial systems have argued that cognitive computations are implemented as structured trajectories and rotations within low-dimensional subspaces embedded in high-dimensional activity space (Chung & Abbott, 2021; Langdon et al., 2023; Vyas et al., 2020). Our findings extended this framework to spatial attention: different cueing paradigms corresponded to distinct rotations on a common cue–choice manifold, and attentional flexibility emerged from dynamically steering a single recurrent circuit through alternative geometric configurations, rather than switching between separate modules.

The same networks that exhibited rich low-dimensional geometry also lacked sharply separated functional clusters for endogenous, exogenous, and social attention. K-means clustering revealed weak cluster structure and highly interleaved unit preferences, indicating that the advanced cognitive module supports all three attention modes within a shared representational substrate. This pattern converged with large-scale neuroimaging work showing that attentional states were more accurately characterized by distributed, dynamically interacting networks rather than isolated, localized foci of activation. Predictive network models have demonstrated that individual differences in sustained and selective attention are captured by whole-brain connectivity patterns spanning dorsal and ventral attention networks, frontoparietal control regions, and default-mode systems, rather than by any single region (Rosenberg et al., 2016, 2017; Wu et al., 2020). Similarly, recent multivariate decoding of feature- and spatial-attention signatures has revealed both shared and distinct large-scale patterns across frontoparietal and occipito-temporal cortex, consistent with partially overlapping attentional codes embedded in a common network architecture (Yang et al., 2025). From a connectome perspective, functional connectome fingerprinting and connectome-based prediction studies further demonstrated that cognitive traits and task activations were reliably encoded in distributed connectivity motifs, not in focal loci (Finn et al., 2015; Tavor et al., 2016). Convergent neuromarker research in affective and pain states also supported this view, identifying condition-specific multivariate patterns that are spatially widespread yet functionally specific (Chang et al., 2015; Krishnan et al., 2016; Zhou et al., 2021). Within this broader network neuroscience framework, the absence of sharply segregated attention-related clusters in our models is better interpreted as a mechanistic realization of distributed coding than as a limitation of the architecture.

Within this distributed substrate, we found that spatial attention was supported by highly sparse activity patterns in the advanced cognitive module. Only a small fraction of recurrent units was required to explain most of the task-related variance and to preserve human-like facilitation–inhibition profiles after virtual lesioning, and the sparsity index approached values reported in primate association cortex. This pattern is consistent with the view that sparse, energy-efficient codes are a general organizing principle of cortical computation. In primary visual cortex, naturalistic stimulation drove activity patterns in which only a minority of neurons responded strongly at any given time, yielding sparse and decorrelated codes that are well suited for efficient sensory representation (Graham & Field, 2007; Vinje & Gallant, 2000). Similar sparse-and-distributed representations have been documented for complex cognitive content such as episodic memories in human hippocampus (Wixted et al., 2014) and in higher-order cortices where dimensionality reduction and concentration of variance onto a limited set of principal axes support robust readout and generalization (Beyeler et al., 2019). Recent fMRI work further suggested that many cognitive states, including working memory and executive control, were expressed through spatially distributed but temporally sparse activation patterns across large-scale networks (Jääskeläinen et al., 2022). Our results extended these ideas to spatial attention: as training progressed, sparsity in the recurrent module increased and strongly predicted reductions in loss, indicating that the network converged toward a compact, low-dimensional representation in which a limited subset of units carried the bulk of attention-relevant information.

Experimental studies provided convergent evidence that such sparse representations were a hallmark of biological attentional control. Single-unit recordings in parietal and frontal areas have shown that only a small proportion of neurons exhibit strong, sustained modulation by covert spatial attention or attentional instructions, while the majority remain weakly tuned or largely unmodulated, resulting in a sparse but behaviorally informative population code (Amengual & Ben Hamed, 2021; Gottlieb et al., 1998). Theoretical and computational work likewise indicated that attention and memory could emerge from recurrent sparse reconstruction or efficient coding mechanisms that selectively amplified a small set of task-relevant features while suppressing redundant ones (Młynarski & Tkačik, 2022; Palm, 2013; Shi et al., 2022). In our ANNs, models endowed with richer visual experience achieved higher sparsity with fewer units and showed the strongest coupling between sparsity and performance, suggesting that experience-dependent optimization push the system toward an efficient regime in which attention is implemented through a small number of strongly tuned, task-critical units embedded in a broader distributed network. Together with the lack of sharply segregated functional clusters, these findings supported a network-level account in which sparse, distributed population codes—rather than dense activity or localized attention modules—provided the neural substrate for flexible endogenous, exogenous, and social orienting.

A further insight was that all three forms of spatial attention relied on strongly mixed-selective codes in the advanced cognitive module, with nonlinear mixed selectivity emerging as the dominant component and tightly tracking the emergence of human-like attention effects. In our models, variance explained by nonlinear conjunctions of cue type, cue direction, and validity grew rapidly during training and plateaued at high levels, whereas the linear component remained comparatively small and slowly varying. At the same time, increases in both the magnitude and heterogeneity of nonlinear mixed selectivity were systematically associated with reductions in training loss across architectures, indicating that conjunctive coding of task variables was not an incidental by-product of learning but a core mechanism supporting flexible attentional behavior. This pattern converged with recent work showing that nonlinear mixed selectivity expanded representational dimensionality and enabled a single population to implement many input–output mappings with simple readouts, thereby supporting context-dependent computation and flexible cognition in high-dimensional state spaces (Ebitz & Hayden, 2021; Fusi et al., 2016; Panzeri et al., 2015; Rigotti et al., 2013; Saxena & Cunningham, 2019). Within this framework, our results suggested that different forms of spatial attention do not recruit isolated endogenous, exogenous, or social-selective units; instead, they are implemented by overlapping ensembles whose nonlinear responses encode specific cue–direction–validity conjunctions, providing a geometric substrate for switching between multiple attentional modes within a shared recurrent circuit.

A broad empirical literature demonstrated mixed selectivity across species, tasks, and brain regions. In macaque prefrontal cortex (PFC), single-unit and population recordings have shown that neurons frequently encode higher-order combinations of task rules, stimuli, and choices during cognitive control, covert attention tasks, working memory, and flexible decision-making, rather than single variables in isolation (Lundqvist et al., 2023; Parthasarathy et al., 2017; Rigotti et al., 2013; Sapountzis et al., 2022; Stokes et al., 2013). Similar conjunctive coding has been observed in mouse PFC during rule-based categorization (Reinert et al., 2021), in primate amygdala and PFC for abstract context and value representations (Saez et al., 2015), and in cortical-hippocampal networks during episodic memory and contextual remapping (Eichenbaum, 2018; McKenzie et al., 2016). Mixed spatial and movement representations have been reported in primate posterior parietal cortex using single-unit recordings (Hadjidimitrakis et al., 2019; Raposo et al., 2014), and partially mixed selectivity has been documented in human parietal association cortex with physiological recordings (Zhang et al., 2017, 2020). These converging results supported a population-based view in which cell assemblies, rather than single feature detectors, formed the fundamental units of cognition (Ebitz & Hayden, 2021; Eichenbaum, 2018; Panzeri et al., 2015; Saxena & Cunningham, 2019; Yuste, 2015). Against this background, the dominance of nonlinear mixed selectivity in our models indicated that human-like spatial attention could be implemented by the same general principle: high-dimensional, mixed-selective population codes that flexibly combine multiple task variables, rather than by segregated channels dedicated to any single attention type.

In conclusion, our findings demonstrated that human-like spatial attention could be realized within a single recurrent circuit, operating in a regime of low-dimensional state-space rotations, sparse population activity, and pervasive nonlinear mixed selectivity. This framework yields concrete, testable predictions for biological systems: changes in attentional demands should systematically reshape the geometry, sparsity, and conjunctive coding of population activity within attentional networks. Future work should focus on extending these models to more naturalistic tasks and incorporating additional biological constraints, such as laminar architecture, neuromodulatory influences, and structural connectivity. Constraining and fitting such networks directly to electrophysiological and neuroimaging data will enable more rigorous ANN–brain comparisons. By jointly aligning behavior, neural geometry, and representational statistics, this approach may help bridge algorithmic and circuit-level accounts of attention, and clarify how distributed, high-dimensional population codes support flexible cognition in the human brain.

## Materials and methods

### Stimuli

Nine visual stimuli were employed: blank, central left/right cues, peripheral left/right cues, social left/right cues, and left/right targets (Fig. 9). The blank stimulus was a uniform gray background (32 × 32 pixels). Central cues were black triangular arrows presented at the center of the gray background, pointing leftward or rightward (Fig. 9a, e). Peripheral cues were black squares placed laterally to the left or right of center (Fig. 9b, f). Social cues depicted eyes with gaze averted leftward or rightward (Fig. 9c, g). Targets were red disks positioned laterally to the left or right of center, identical across conditions (Fig. 9d, h). All stimuli followed established spatial cueing paradigms in human studies, with the only modification being color differentiation between cues (black) and targets (red).

**Fig. 9.**
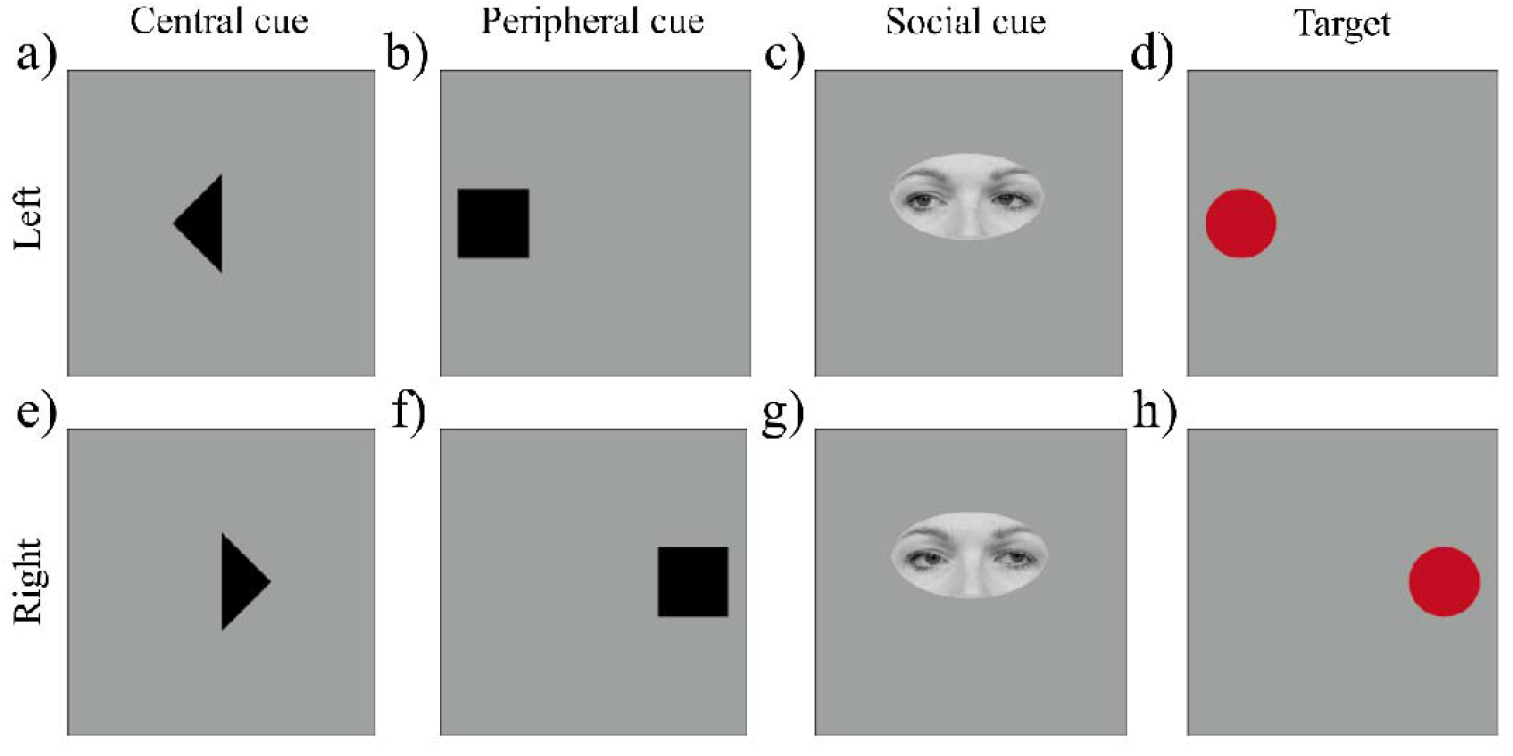
Visual stimuli used in the spatial cueing tasks. (a) Left-pointing central cue; (b) Left peripheral cue; (c) Left social cue; (d) Left target; (e) Right-pointing central cue; (f) Right peripheral cue; (g) Right social cue; (h) Right target.

### Experimental Tasks

The experiment comprised three cueing paradigms, distinguished by cue type: (1) central cueing, (2) peripheral cueing, and (3) social cueing. Central cueing, designed to engage endogenous attention, utilized a CV of 70%, consistent with prior findings that robust cueing effects emerge when cues are predictive (Riggio & Kirsner, 1997). Peripheral cueing, primarily engage exogenous attention, employed a 50% CV, rendering cues non-predictive and thereby minimizing endogenous contributions while maximizing reflexive orienting (Posner & Cohen, 1984). Social cueing, also set at 50% CV, probed attention orienting elicited by gaze stimuli. RTs were generated across seven different CTOAs (100, 250, 300, 500, 800, 1200 and 2400ms) to characterize temporal dynamics of spatial selective attention. RT data were modeled on human performance reported in prior studies for central (Cheal & Lyon, 1991; Muller & Rabbitt, 1989; Posner & Cohen, 1984), peripheral (Cheal & Lyon, 1991; Klein, 2000; Mulckhuyse & Theeuwes, 2010; Posner & Cohen, 1984; Samuel & Kat, 2003; Song et al., 2014; Vossel, Mathys, et al., 2014), and social cueing (Driver et al., 1999; Friesen & Kingstone, 1998, 2003; Frischen & Tipper, 2004; Langton & Bruce, 1999; Mulckhuyse & Theeuwes, 2010; Xu et al., 2011). Full parameter details are listed in Table S2.

Each trial consisted of five sequential periods: (1) an initial blank screen (200ms) serving as a preparatory period; (2) cue presentation, during which a central, peripheral, or social cue indicated a potential target location (left/right), with durations specified in Table S3; (3) a delay interval (blank screen) separating cue offset and target onset, varying by CTOA (see Table S3); (4) target presentation, a red disk displayed for 100ms at one of the target location; (5) a post-target blank screen (200ms) providing the response window. RTs, fixed based on previous empirical studies to ensure consistency with established attention effects, were detailed in Table S2. For analysis, to improve training stability, RTs were converted to reaction speed (RS = 1/RT) and normalized to the range [0, 1] using min-max scaling:

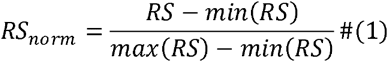

where 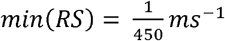, and 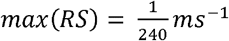. To balance conditions, 840 trials were generated (3 cue types × 7 CTOAs × 40 repetitions). Within each CTOA, trial distributions across target locations (left/right) and validity (valid/invalid) followed the CV of each paradigm (Table S4).

### ANN structure

The ANN model was designed as a computational model to simulate human performance across three visual attention tasks — endogenous, exogenous, and social attention. The network received stimulus images (central, peripheral, and social cues, blank screens, and targets) as inputs and generated behavioral responses (i.e., RS) as outputs, The ANN architecture comprised three modules (Fig. 1b): (1) a sensory module for visual feature extraction; (2) a cognitive module for attentional processing; and (3) an action module for response generation.

The sensory module, implemented as a CNN, extracted high-level visual features from input stimuli. A Pretrained AlexNet, trained on the CIFAR-10 natural image dataset, was used to endow the model with basic visual processing capabilities. During subsequent training on attention tasks, the sensory module weights were frozen, and only the cognitive and action modules were updated.

The cognitive module, representing higher-level cognitive processes, was implemented as a recurrent natural network (RNN) with *N*_*rec*_ = 256 leaky recurrent The cognitive module, representing higher-level cognitive processes, was units. Its dynamics followed:

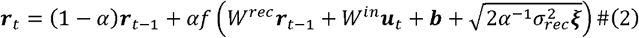

where ***r***_*t*_ and ***u***_*t*_ represent recurrent activations and inputs from the sensory module at time *t*, respectively. *W*_*in*_ and *W*_*rec*_ separately denote input and recurrent weight matrices, ***b*** the bias, and *ξ* Gaussian noise (mean = 0, standard deviation = 1) with strength *σ*_*rec*_ = 0.01 The parameter *α* ≡ Δ*t* / *τ* with Δ*t*, = 10ms as the time step and *τ* = 100ms as the membrane constant. The activation function was rectified linear (ReLU).

The action module mapped recurrent activity to motor outputs via:

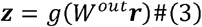

where ***z*** is the output vector, *W*^*out*^ the output weight matrix, and *g* (*x*) a sigmoid function constraining ***z*** to [0, 1]. The module consisted of three output units: a fixation unit and two response units (left, right). The fixation unit was active (output = 1) before target onset and deactivated (output = 0) during target and post-target periods. Upon target onset, the corresponding response unit was activated while the other remained silent.

The normalized response speed (*RS*_*norm*_) was defined as the maximum activation of the response units during post-target period:

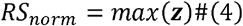

### Sensory module variations

To examine how sensory architecture and pretraining influence attentional performance, we compared the Pretrained AlexNet model (described above) with the following five variants (Fig. 2):

1. Pretrained ResNet: similar to the Pretrained AlexNet, but employing a CIFAR-10 natural image pretrained ResNet18 (He et al., 2016) as the sensory module.
2. Stimulus AlexNet: the sensory module followed the AlexNet architecture but was pretrained solely on experimental stimuli before attention task training, providing task-specific visual experience. Its sensory weights were frozen thereafter, with only the cognitive and action modules updated.
3. End-to-end AlexNet: the AlexNet architecture was used without pretraining. All parameters across sensory, cognitive, and action modules were trained jointly, allowing sensory representations to emerge alongside task-specific learning.
4. Random AlexNet: the sensory module was a randomly initialized AlexNet with fixed weights throughout, leaving only the cognitive and action modules trainable.
5. Random ResNet: similar to the Random AlexNet, but employing a randomly initialized ResNet18 (He et al., 2016) as the sensory module with fixed weights.

In summary, the Pretrained AlexNet, Pretrained ResNet, and Stimulus AlexNet incorporated prior visual experience (general vs. task-specific), the End-to-end AlexNet acquired visual experience during attention task training, whereas the Random AlexNet and Random ResNet lacked pretraining and fixed their sensory weights, providing no learned visual representations.

### Network training procedure

Models were trained using backpropagation through time to minimize a mean squared error (MSE) loss function:

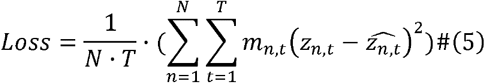

where *n* and *t* index units and timesteps, respectively, *z*_*n,t*_ denotes the target output, and 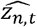 the model prediction. The masking term *m*_*i,t*_ was set to 0 during the 100 ms target presentation (serving as a grace period), to 50 during the post-target period to accelerate learning of the correct response, and to 1 otherwise.

Training employed the Adam optimizer (Kingma & Ba, 2017) with a learning rate of 0.001, batch size of 128, for 1400 epochs. To assess training stability, we trained 50 independent instances of the Pretrained AlexNet model with identical model hyperparameters, initialization procedures, and training parameters.

### mTDR analysis

We applied the mTDR method to characterize population activity, decomposing the cognitive module’s trial-by-trial neural responses into task-variable-specific, low-dimensional encoding subspaces (Aoi et al., 2020). Units with low maximum activation (< 0.1) were removed. At the single-unit level, the activity of unit on *n* on trial *k* at time *t, y*_*k,t,i*_, was modeled as a linear combination of *P* task variables 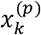:

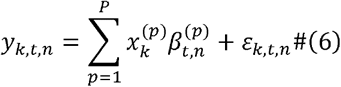

Where *P* is 3 (i.e., 3 task variables: cue, choice, bias), cue variable 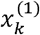 takes on values of ±1, indicating a left or right cue, and choice variable 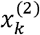 also takes on values of ±1, indicating a left or right choice. 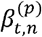 are time-varying unknown weights. *ε*_*k,t,n*_ is noise.

Concatenating responses of all *N* units into a vector ***y***_*k,t*_, and across all ***T*** time points on trial *k*, formed the observation matrix ***Y***_*k*_ = [ ***y***_*k*,1_, ***y***_*k*,2,_ …, ***y***_*k,T*_] *Y*_*k*_ could be decomposed as follows:

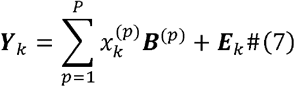

where ***B***^(*p*)^ denotes a *N* units × *T* time matrix of unknown weights for the *p*^*th*^ task variable: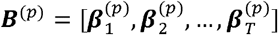, and 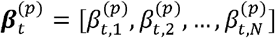. ***E***_*k*_ is a *N* units × *T* time matrix of noise: ***E***_*k*_ = [ ***ε***_*k*,1_, ***ε***_*k*,2,_ …, ***ε***_*k,T*_] and, ***ε***_*k,t*_ = [ *ε*_*k,t*,1,_ *ε*_*k,t*,2,_ … *ε*_*k,t,N*_].

To obtain a compact description we factorized each ***B*** ^(*p*)^ with low-rank factorization ***B***^(*p*)^ = ***W***^(*p*)^ ***S***^(*p*)^,where ***W*** ^(*p*)^is a *N* × *r* ^(*p*)^ matrix of unit-dependent mixing weights and ***S***^(*p*)^ is a *r* ^(*p*)^ × *T* matrix of time-varying basis functions. *r* ^(*p*)^ = *rank* (***B*** ^(*p*)^), namely, *r* ^(*p*)^ is the dimensionality of the encoding of the *p*^*th*^ task variable. The equation (7) thus became:

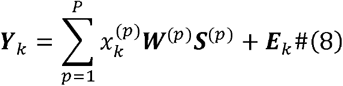

The ranks *r* ^(*p*)^ were chosen using a stepwise Akaike information criterion (AIC): additional dimensions were included only when they produced a lower AIC value. To obtain a unique and interpretable set of encoding axes for visualization, we orthogonalized the subspaces of ***W***^(*p*)^sequentially by QR decomposition process. For each cue type and CTOA condition, trial-averaged population vectors 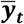 were then projected into each encoding subspace as the following equation to yield low-dimensional trajectory:

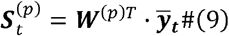

We further visualized neural population geometry by projecting each trial onto a two-dimensional space defined by the cue axis (horizontal) and the choice axis (vertical). Rightward and leftward cues map to the right and left sides of the cue axis, respectively, whereas rightward and leftward choices map to the upper and lower sides of the choice axis. A population representation trajectory aligned with the vertical choice axis indicates target encoding independent of validity, suggesting the absence of attentional modulation. By contrast, trajectories falling in the first and third quadrants reflect facilitation, with neural activity biased toward cued locations, whereas trajectories in the second and fourth quadrants reflect inhibition, with encoding biased toward uncued locations. This quadrant-based framework thus provides an intuitive geometric readout of whether, at a given CTOA and for each cue type, neural population representations favor facilitation or inhibition. To quantify these effects, we computed the acute angle θ between the symmetry axis of cue-specific trajectories and the vertical choice axis. Values of θ near zero indicate no systematic cue modulation, whereas positive (clockwise) and negative (counter-clockwise) deviations correspond to increasing facilitation and inhibition, respectively.

### Functional cluster analysis

To investigate the functional organization of neural activity within the cognitive module, we applied k-means clustering to activation pattern. For each cue type, three types of activation features were extracted: (1) mean activation during the cue period for valid and invalid conditions separately; (2) mean activation during the target-to-trial-end for valid and invalid conditions separately; and (3) attention effects, defined as the difference in mean activation between valid and invalid conditions across the seven CTOAs. Thus, each unit was characterized by 33 features in total (2 periods × 2 validity × 3 cue types + 7 CTOAs × 3 cue types).

All features were z-scored across conditions to emphasize unit’s relative selectivity, and each unit’s resulting feature vector was used as input to k-means clustering.

Cluster quality was evaluated using the Silhouette score (G. R. Yang et al., 2019):

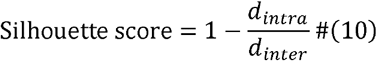

where *d*_*intra*_ is the mean intra-cluster distance for a unit, and *d*_*inter*_ is its mean distance to units in other clusters. The Silhouette score ranges from 0 to 1, with higher values indicating more distinct cluster separation.

### Ablation analysis of units in the cognitive module

To estimate the minimal subset of units required to capture task-related variability and support behavioral performance, we conducted an ablation analysis within the cognitive module. For each unit, mean activations were calculated across trial time points (excluding the initial blank) for all 42 conditions (3 cue types × 7 CTOAs × 2 cue valid/invalid). The variance of these condition-specific means was then computed. Variance values were normalized by the total variance across all units, constraining the cumulative variance explained sum to 1, and ranked in descending order, yielding a cumulative variance explained curve that provided an estimate of the number of units sufficient to account for representation variability of the module.

We then assessed functional contribution to behavior through progressive unit ablation. Units were eliminated sequentially, beginning with those explaining the greatest variance; for each step, both the selected unit and all units with lower variance were lesioned by setting their input and output connection weights to zero. The network was subsequently evaluated on the same attention tasks, and the change in *Loss* relative to the intact model served as a quantitative measure of the unit’s contribution. This stepwise elimination provided a lower-bound estimate of how many units necessary to preserve task-relevant performance.

### Sparsity analysis

We quantified representational sparsity in the cognitive module, using the sparsity index proposed by Vinje & Gallant (2000):

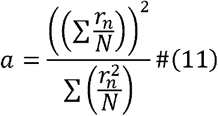

where *r*_*n*_ is the activation of unit *n*, and *N* is the total number of units. This value was standardized to yield the sparsity measure *S*:

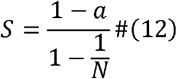

with *S* ranging from 0 to 1, where higher values indicate greater sparsity.

During training we sampled every 10 epochs, and extracted (i) model performance, measured as training loss (*Loss*) and (ii) sparsity *S*. To assess the relationship between sparsity and performance, we fitted a linear regression across epochs:

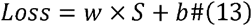

where the slope *w* indexes the impact of sparsity on performance and *b* is the intercept. A positive *w* indicated that increased sparsity predicted higher loss (worse performance), whereas a negative *w* indicated that increased sparsity predicted improved performance. The two parameters were estimated by ordinary least squares.

### Mixed selectivity analysis

To characterize how units within the cognitive module encode combinations of task variables — cue type (central (C), peripheral (P) and social (S)), cue direction (left (L) and right (R)), and validity (invalid (IV) and valid (V)) — we adopted the mixed selectivity framework (Rigotti et al., 2013), decomposing each unit’s response into pure, linear mixed, and nonlinear mixed selectivity components. Units with low overall activation values (maximum activation < 0.1) were excluded.

For each unit *n* within a specific time bin, we fitted a three-factor regression model to capture linear selectivity components of its activation *r*_*n*_:

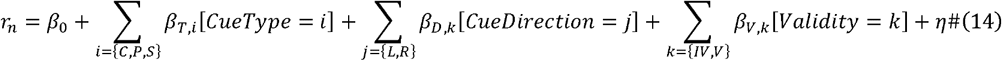

where [ ·] denotes task variables, *β*_0_ is a bias term, and η is Gaussian noise. Units were classified as pure selective if only one main effect coefficient was significant (p < 0.05), and as linear mixed selective if multiple main effects were significant but no interaction terms improved the model fit.

To capture nonlinear mixed selectivity component, we computed residual activations not explained by the above linear model:

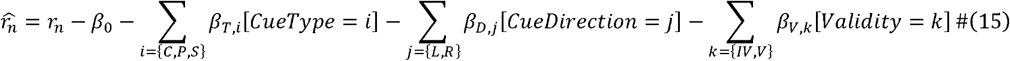

and then fitted a one-factor interaction model:

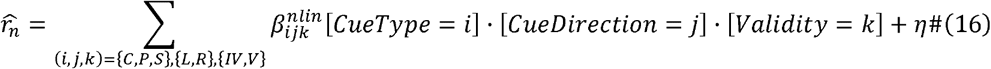

Units with significant interaction coefficients (p < 0.05) were classified as nonlinear mixed selective. Such units enriched representational dimensionality and supported flexible decoding.

At the population level, we quantified linear and nonlinear mixed selectivity by computing the variance in activity explained by each component. For unit *n*, we defined:

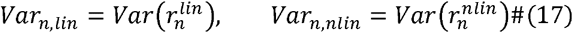

where 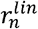 and 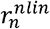 denote activation of the unit *n* explained by the linear and nonlinear components, respectively. Mean variance across the neural population was then computed as

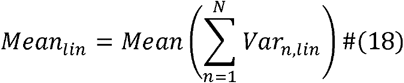

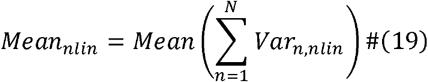

where *N* is the total number of units. In addition, for each component we estimated 95% confidence intervals (CIs) of *Var*_*n,lin*_ and *Var*_*n,nlin*_ across units, and quantified their widths, *ΔCI*_*lin*_ and *ΔCI*_*nlin*_, as the difference between the upper and lower CI bound, providing a summary of across-unit variability (i.e., diversity) in mixed selectivity.

To evaluate their functional relevance of mixed selectivity, we extracted cognitive module activity and training loss (*Loss*) every 10 epochs to fit the following linear regressions:

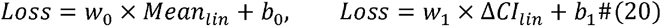

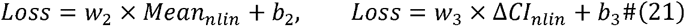

where slopes *w*_0_ and *w*_1_ separately index the impact of the mean and dispersion of linear mixed selectivity on model performance, and *w*_2_ and *w*_3_ play the analogous roles for the nonlinear component. *b*_0_, *b*_1_, *b*_2_, and *b*_3_ are the intercepts. A negative slope indicates that stronger or more heterogeneous mixed selectivity predicts better performance (lower loss), whereas a positive slope indicates the opposite. All parameters were estimated using ordinary least squares.

To further summarize the balance between linear and nonlinear tuning at the population level (restricted to units showing any mixed selectivity), we defined for each unit *n*, a population-level mixed-selectivity preference, *I*_*n,mixed*_ (adapted from Rigotti et al., 2013):

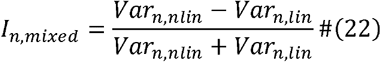

This index is bounded between −1 and 1. Positive values indicate that the unit is more strongly driven by nonlinear (conjunctive) interactions than by linear terms, whereas negative values denote predominantly linear mixed selectivity.

## Code and data availability

All analysis and model code are publicly available at https://github.com/ChenLumen/attention_rnn. All data are publicly available at https://huggingface.co/datasets/mcliu/attention_rnn.

## Supplementary Materials

**Table S1.**
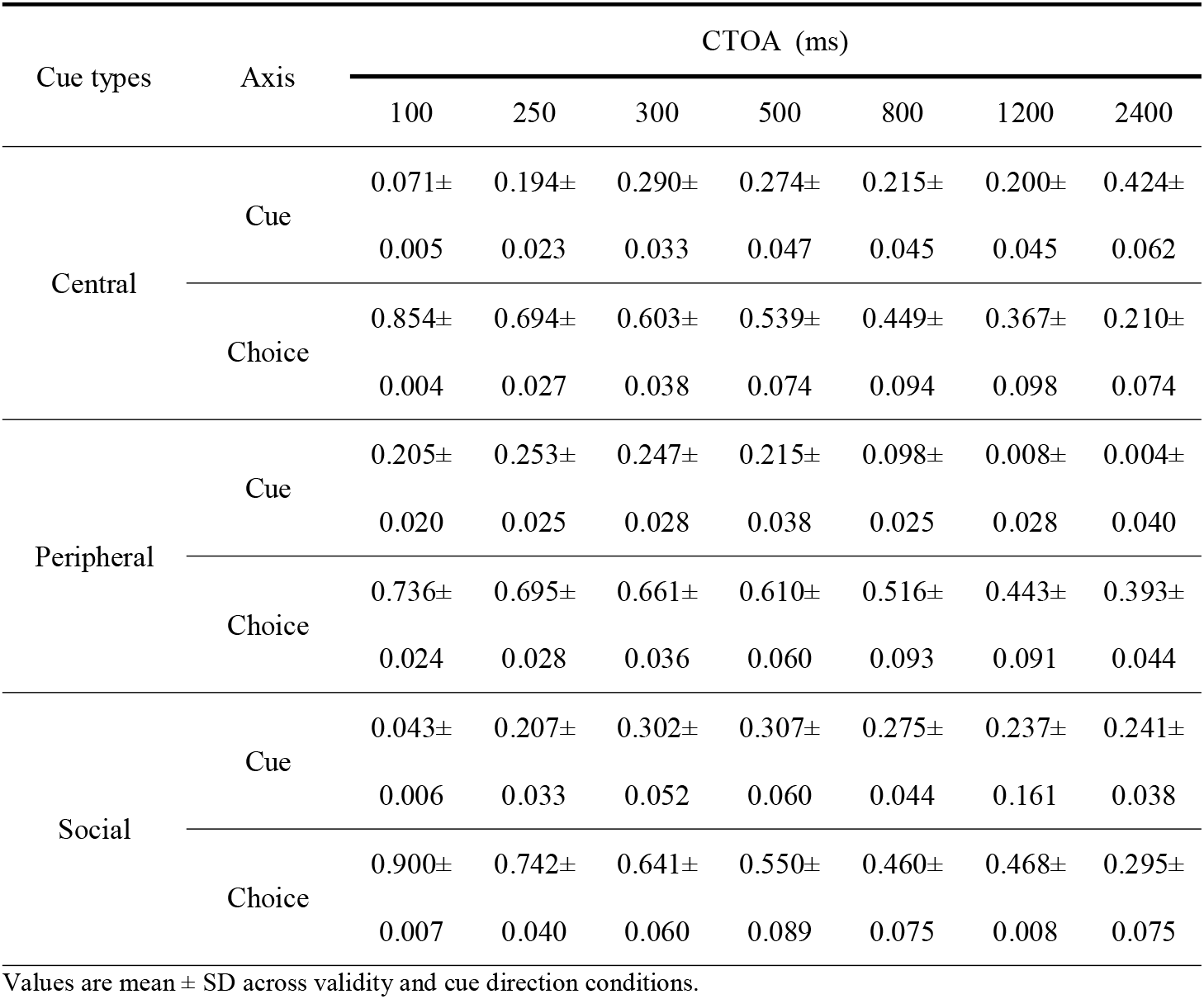
Variance explained (*R*^2^) by the cue and choice axes in the mTDR analysis for the Pretrained AlexNet model across cue types and CTOAs.

| Cue types | Axis | CTOA (ms) |  |  |  |  |  |  |
| --- | --- | --- | --- | --- | --- | --- | --- | --- |
|  |  | 100 | 250 | 300 | 500 | 800 | 1200 | 2400 |
| Central | Cue | 0.071± | 0.194± | 0.290± | 0.274± | 0.215± | 0.200± | 0.424± |
|  |  | 0.005 | 0.023 | 0.033 | 0.047 | 0.045 | 0.045 | 0.062 |
|  | Choice | 0.854± | 0.694± | 0.603± | 0.539± | 0.449± | 0.367± | 0.210± |
|  |  | 0.004 | 0.027 | 0.038 | 0.074 | 0.094 | 0.098 | 0.074 |
| Peripheral | Cue | 0.205± | 0.253± | 0.247± | 0.215± | 0.098± | 0.008± | 0.004± |
|  |  | 0.020 | 0.025 | 0.028 | 0.038 | 0.025 | 0.028 | 0.040 |
|  | Choice | 0.736± | 0.695± | 0.661± | 0.610± | 0.516± | 0.443± | 0.393± |
|  |  | 0.024 | 0.028 | 0.036 | 0.060 | 0.093 | 0.091 | 0.044 |
| Social | Cue | 0.043± | 0.207± | 0.302± | 0.307± | 0.275± | 0.237± | 0.241± |
|  |  | 0.006 | 0.033 | 0.052 | 0.060 | 0.044 | 0.161 | 0.038 |
|  | Choice | 0.900± | 0.742± | 0.641± | 0.550± | 0.460± | 0.468± | 0.295± |
|  |  | 0.007 | 0.040 | 0.060 | 0.089 | 0.075 | 0.008 | 0.075 |

Values are mean ± SD across validity and cue direction conditions.

**Table S2.** Mean RTs for valid and invalid trials across the three spatial cueing paradigms and CTOAs.

| RT (ms) |  | CTOA (ms) |  |  |  |  |  |  |
| --- | --- | --- | --- | --- | --- | --- | --- | --- |
|  |  | 100 | 250 | 300 | 500 | 800 | 1200 | 2400 |
| Central | Invalid | 300 | 297 | 296 | 282 | 276 | 271 | 269 |
|  | Valid | 290 | 282 | 280 | 266 | 260 | 255 | 253 |
| Peripheral | Invalid | 307 | 290 | 280 | 268 | 260 | 257 | 255 |
|  | Valid | 290 | 290 | 288 | 282 | 277 | 271 | 261 |
| Social | Invalid | 298 | 298 | 296 | 282 | 274 | 260 | 250 |
|  | Valid | 290 | 293 | 280 | 266 | 264 | 260 | 260 |

**Table S3.** Cue and delay durations across the three spatial cueing paradigms and CTOAs.

| Cue types | Trial periods | CTOA (ms) |  |  |  |  |  |  |
| --- | --- | --- | --- | --- | --- | --- | --- | --- |
|  |  | 100 | 250 | 300 | 500 | 800 | 1200 | 2400 |
| Central | Cue duration | 100 | 150 | 200 | 200 | 200 | 200 | 200 |
|  | Delay duration | 0 | 100 | 100 | 300 | 600 | 1000 | 2200 |
| Peripheral | Cue duration | 100 | 100 | 100 | 100 | 100 | 100 | 100 |
|  | Delay duration | 0 | 150 | 200 | 400 | 700 | 1100 | 2300 |
| Social | Cue duration | 100 | 150 | 200 | 200 | 200 | 200 | 200 |
|  | Delay duration | 0 | 100 | 100 | 300 | 600 | 1000 | 2200 |

**Table S4.** Trial numbers for each cue type, cue location, and target location across the three spatial cueing tasks at each CTOA.

| Cue types | Cue direction |  | Target location |
| --- | --- | --- | --- |
|  | Left | Right |  |
| Central | 14 | 6 | Left |
|  | 6 | 14 | Right |
| Peripheral | 10 | 10 | Left |
|  | 10 | 10 | Right |
| Social | 10 | 10 | Left |
|  | 10 | 10 | Right |

**Fig. S1.**
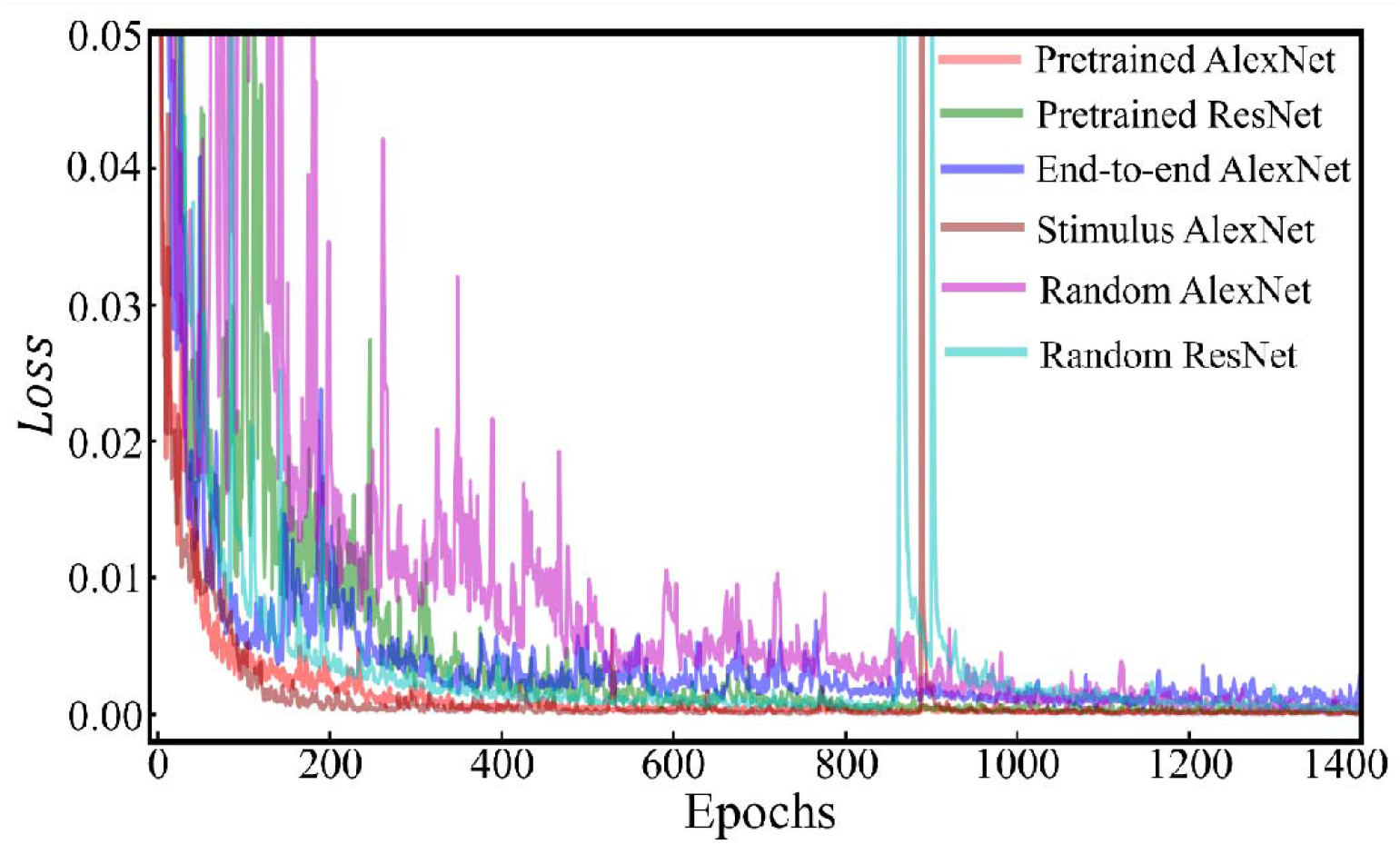
Learning curve of all six models (Pretrained AlexNet, Pretrained ResNet, End-to-end AlexNet, Stimulus AlexNet, Random AlexNet, Random ResNet).

**Fig. S2.**
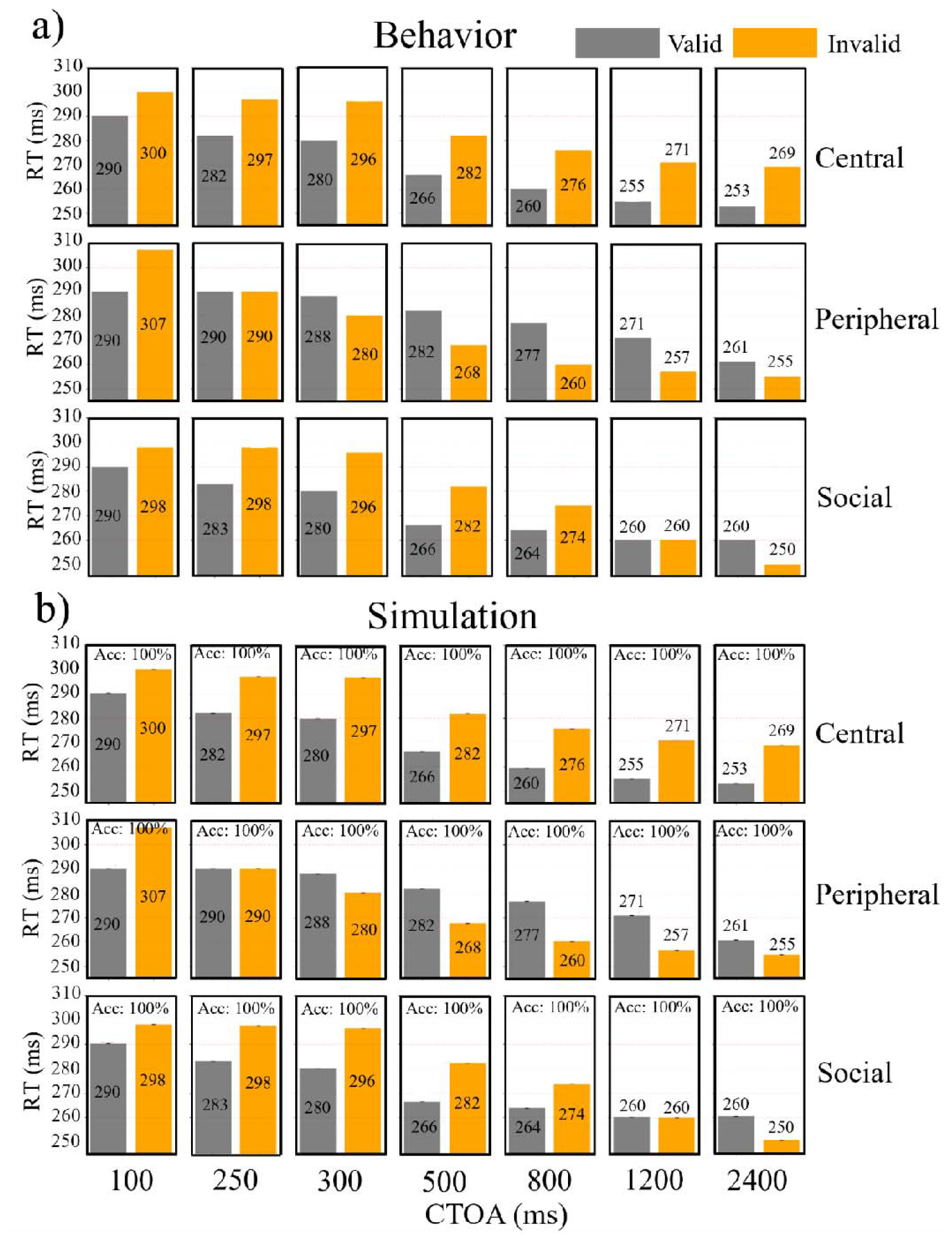
RTs for valid and invalid cues in behavior and the Pretrained AlexNet. (a) Human RTs for valid (gray) and invalid (orange) cues in central, peripheral, and social cueing tasks across CTOAs. (b) RTs produced by the Pretrained AlexNet model for the same three attention tasks and CTOAs.

**Fig. S3.**
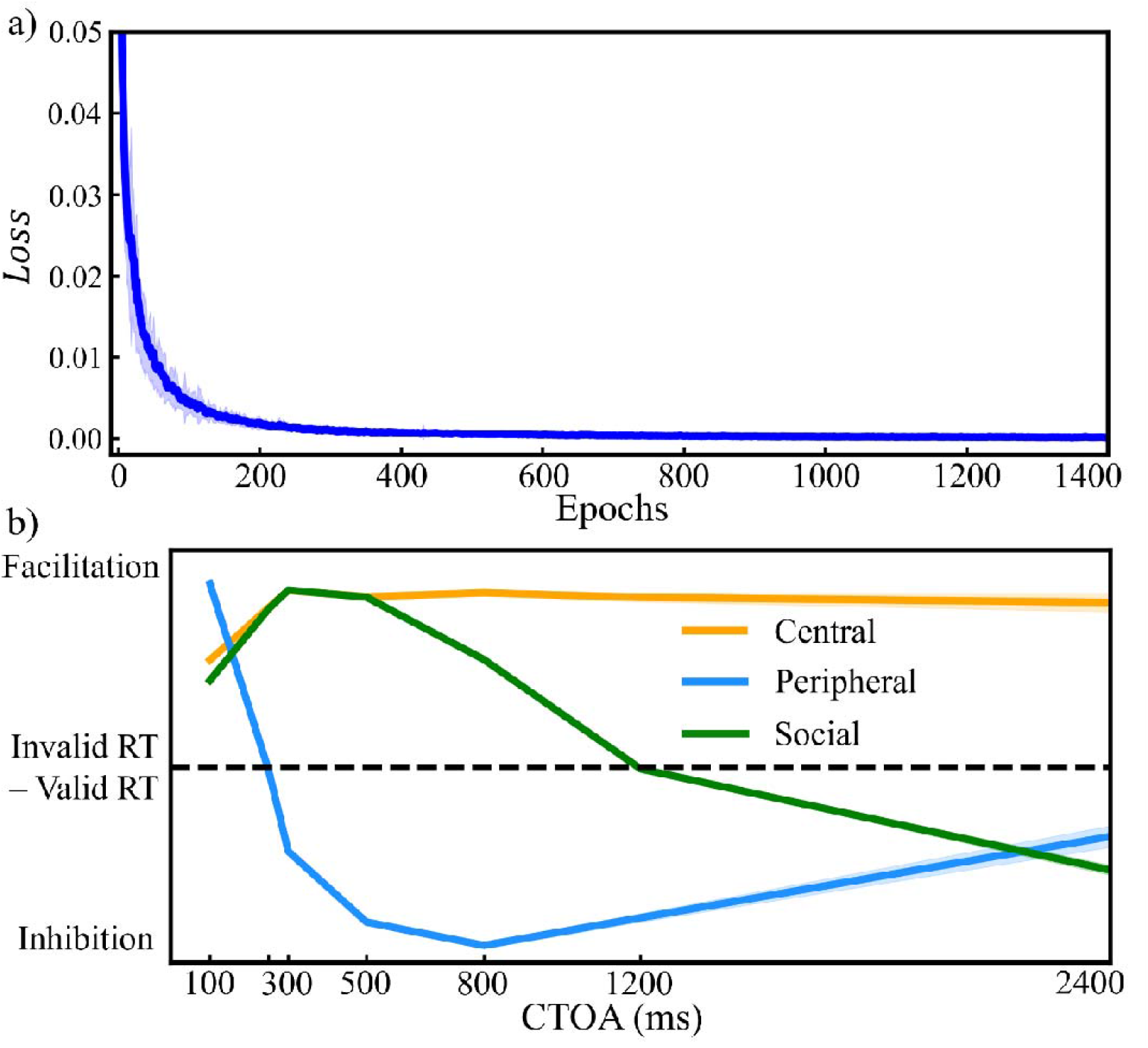
Training stability and behavioral fit of the Pretrained AlexNet model. Fifty networks with identical architecture and hyperparameters were trained independently. (a) Training loss as a function of epochs. (b) Simulated attention effects for central, peripheral, and social cueing across CTOAs. Solid lines indicate the across-network mean, and shaded regions show ±1 standard deviation around the mean.

**Fig. S4.**
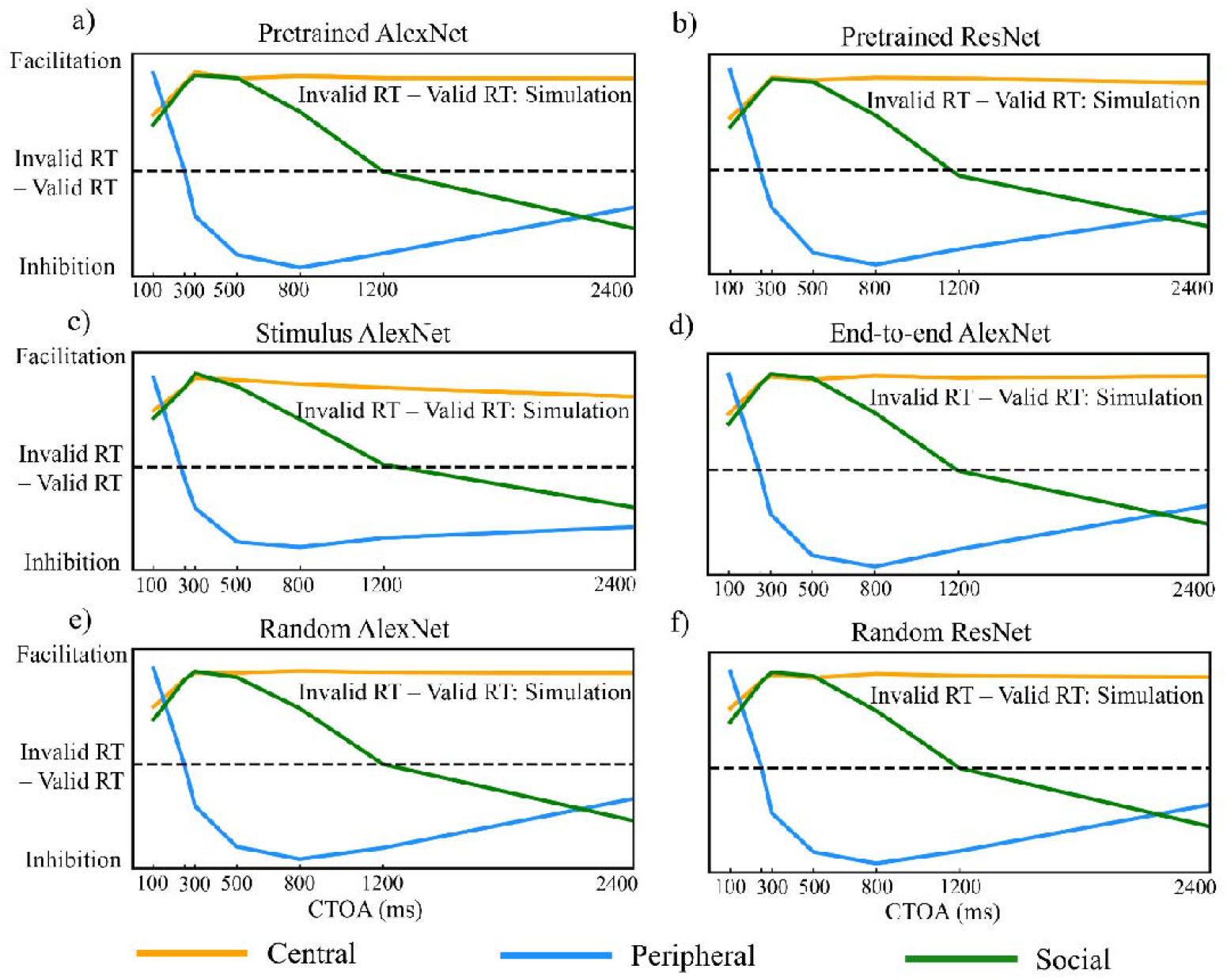
Simulated attention effects across network variants. (a) Pretrained AlexNet; (b) Pretrained ResNet; (c) Stimulus AlexNet; (d) End-to-end AlexNet; (e) Random AlexNet; (f) Random ResNet.

**Fig. S5.**
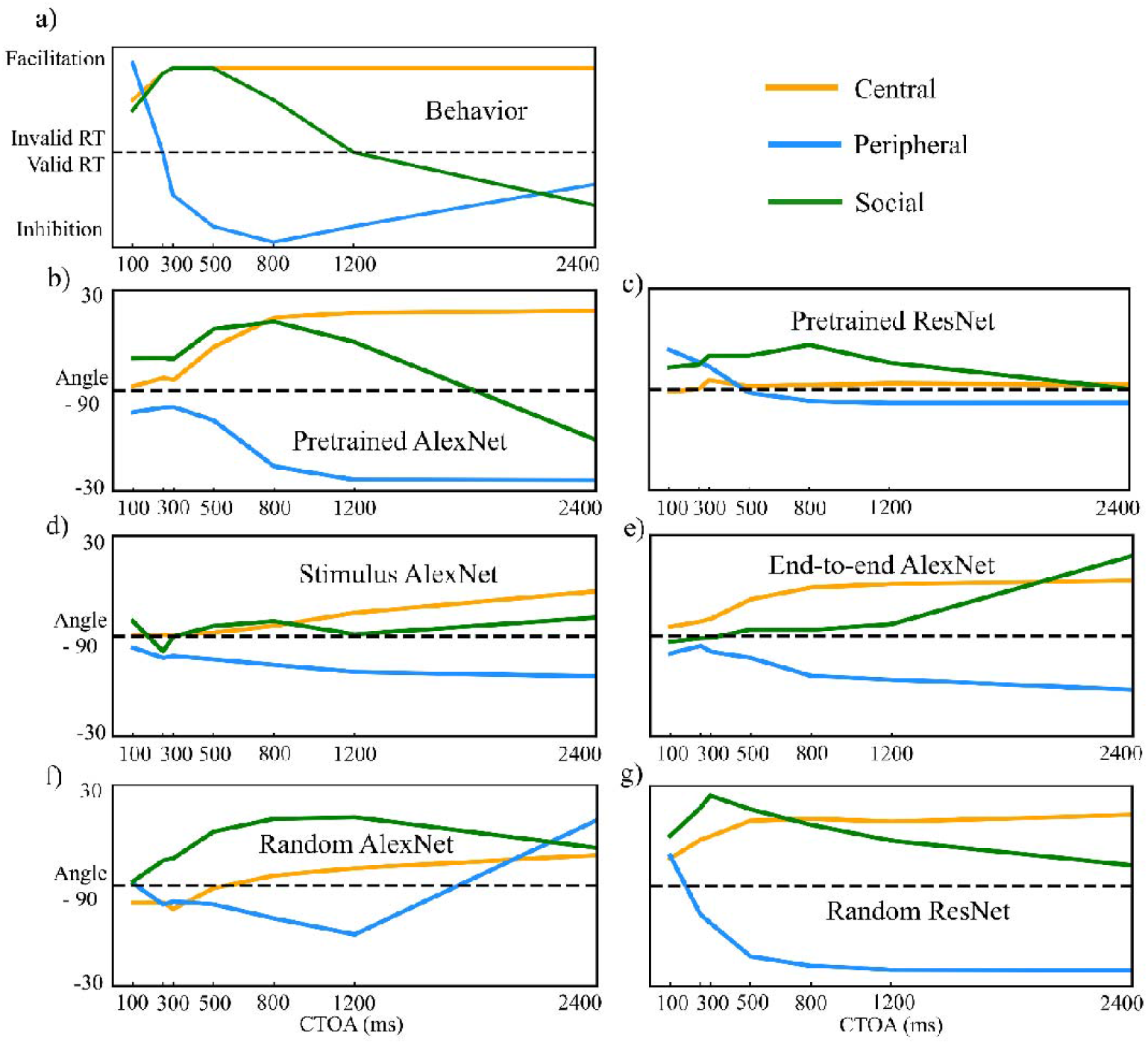
Behavioral attention effects and neural geometry across network variants. (a) Human spatial attention effects, expressed as invalid–minus–valid RTs differences for central, peripheral, and social cueing paradigms. (b–g) Deflection angles of cue-related population trajectories relative to the neutral axis (90°), obtained from the mTDR analysis as a function of CTOA for each model: (b) Pretrained AlexNet; (c) Pretrained ResNet; (d) Stimulus AlexNet; (e) End-to-end AlexNet; (f) Random AlexNet; (g) Random ResNet.

**Fig. S6.**
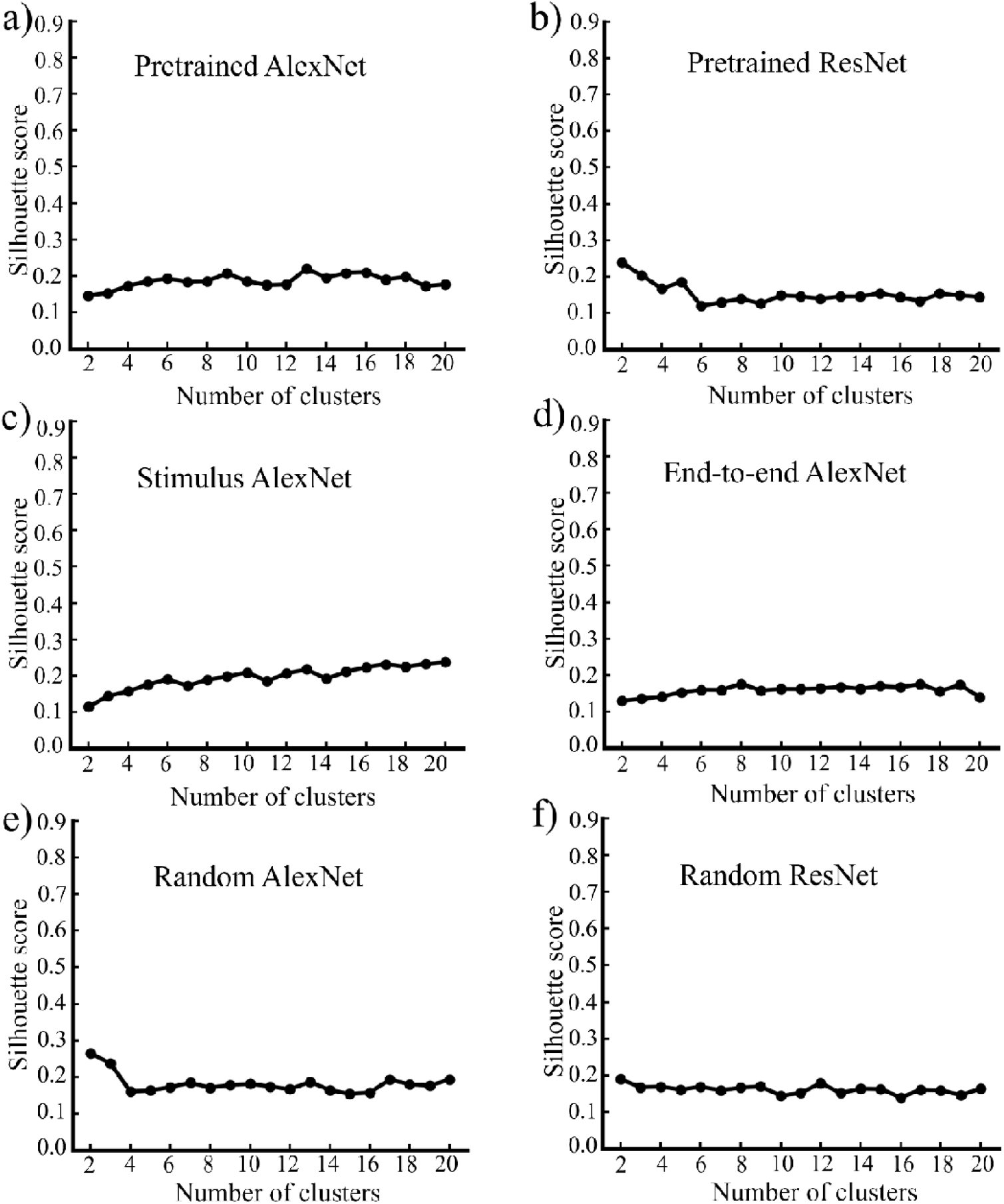
Silhouette scores for functional clustering across network variants. (a) Pretrained AlexNet; (b) Pretrained ResNet; (c) Stimulus AlexNet; (d) End-to-end AlexNet; (e) Random AlexNet; (f) Random ResNet.

**Fig. S7.**
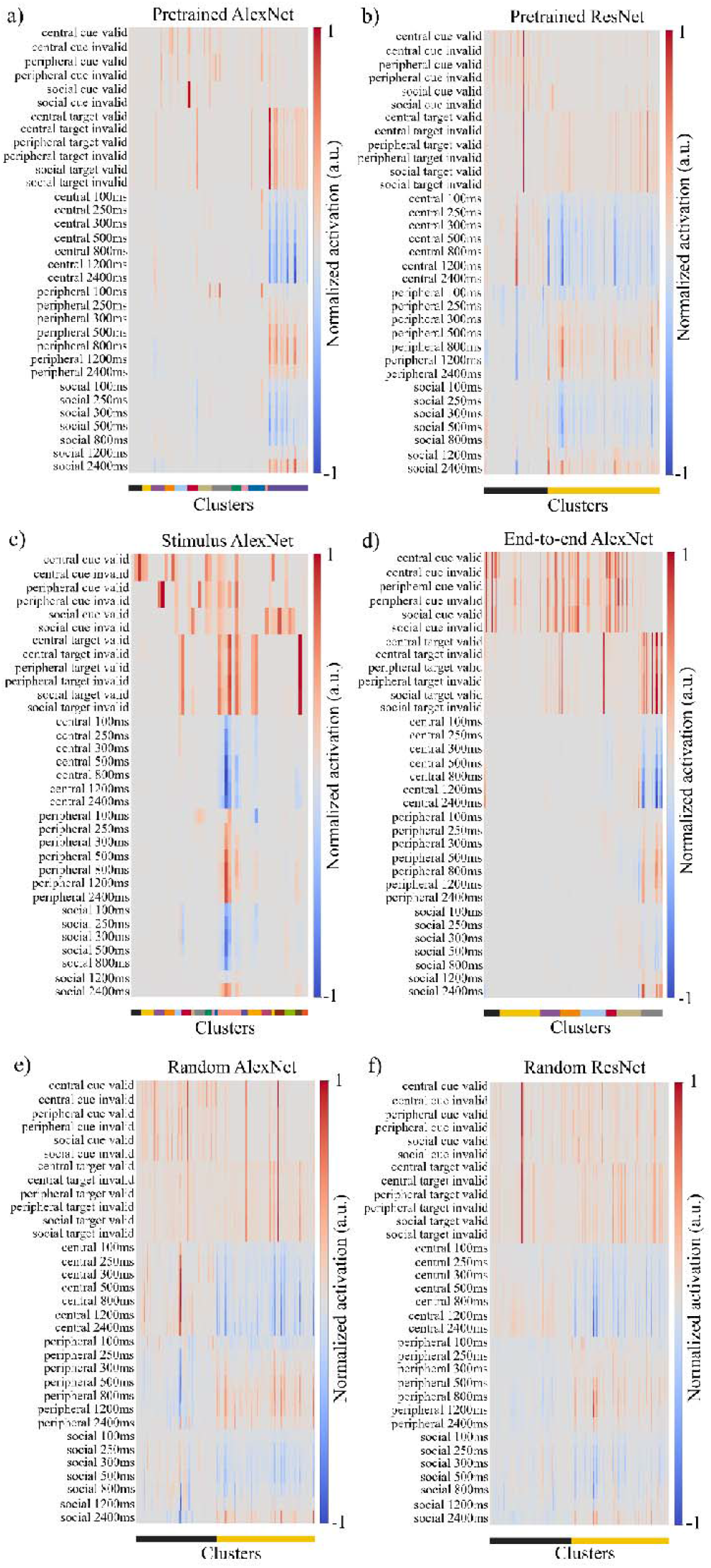
Functional clustering across network variants. (a) Pretrained AlexNet; (b) Pretrained ResNet; (c) Stimulus AlexNet; (d) End-to-end AlexNet; (e) Random AlexNet; (f) Random ResNet. The colored bar beneath each panel indicated cluster identity.

**Fig. S8.**
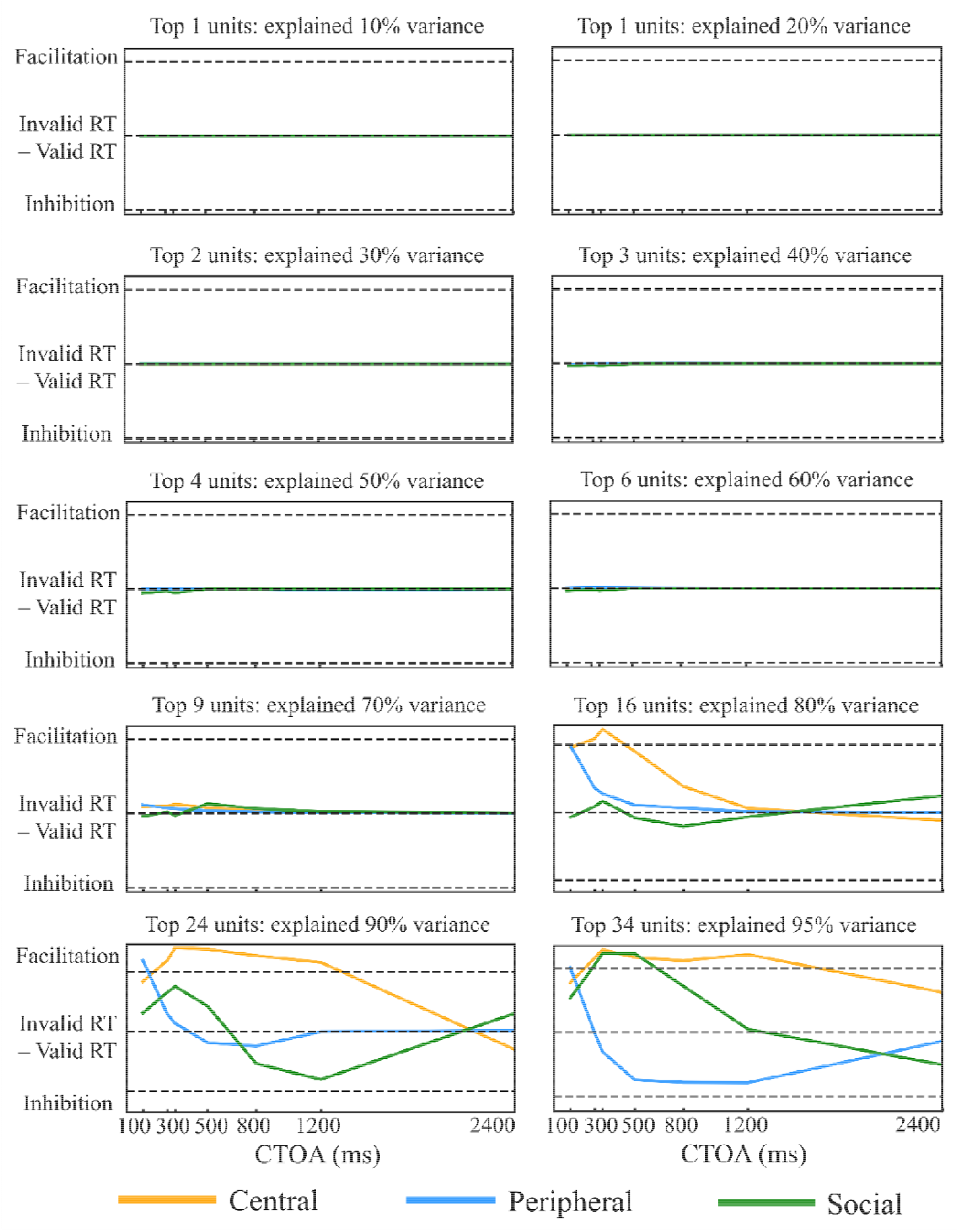
Simulated spatial-attention effects as a function of the proportion of variance retained in the Pretrained AlexNet model.

**Fig. S9.**
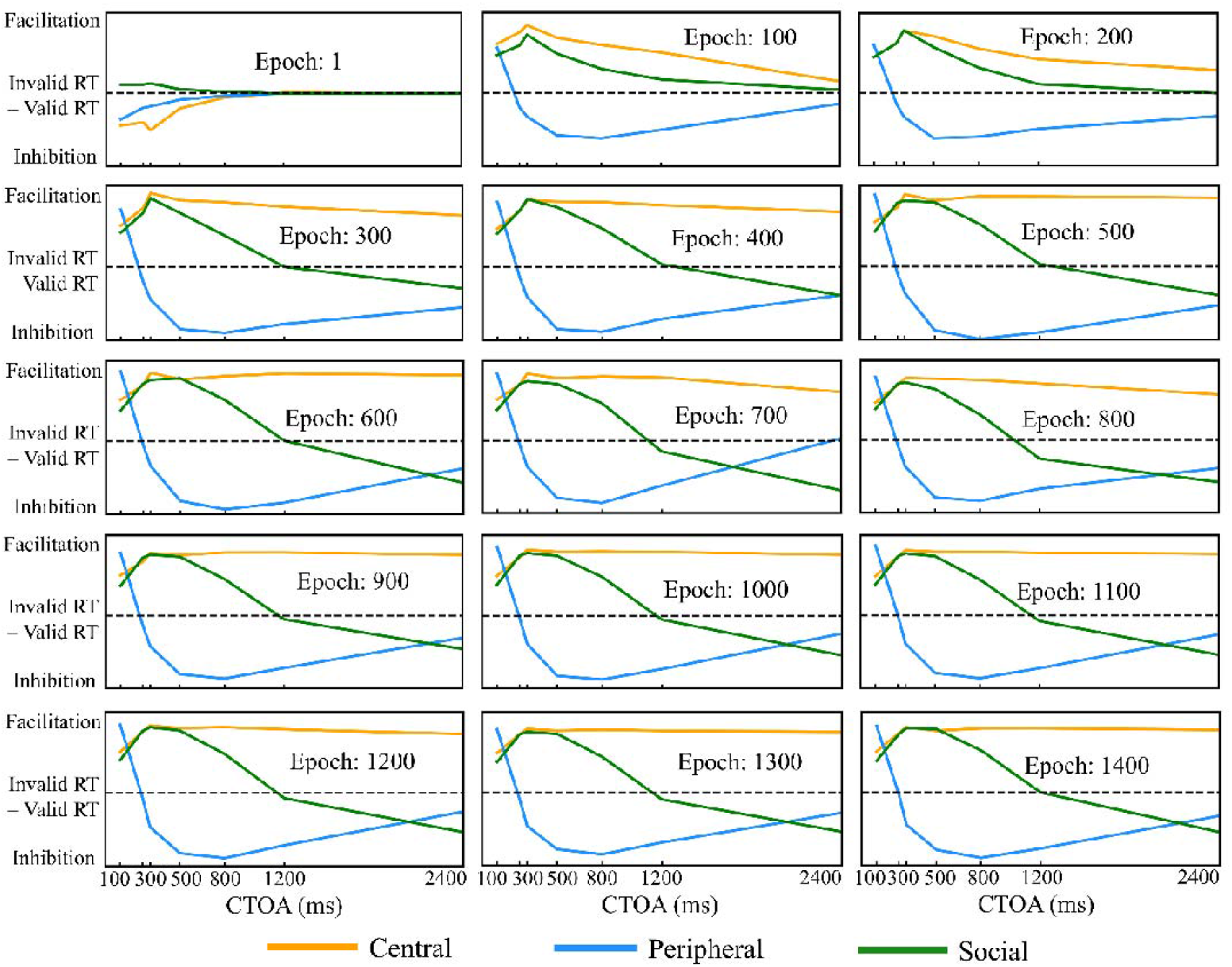
Evolution of simulated spatial-attention effects during training in the Pretrained AlexNet model.

**Fig. S10.**
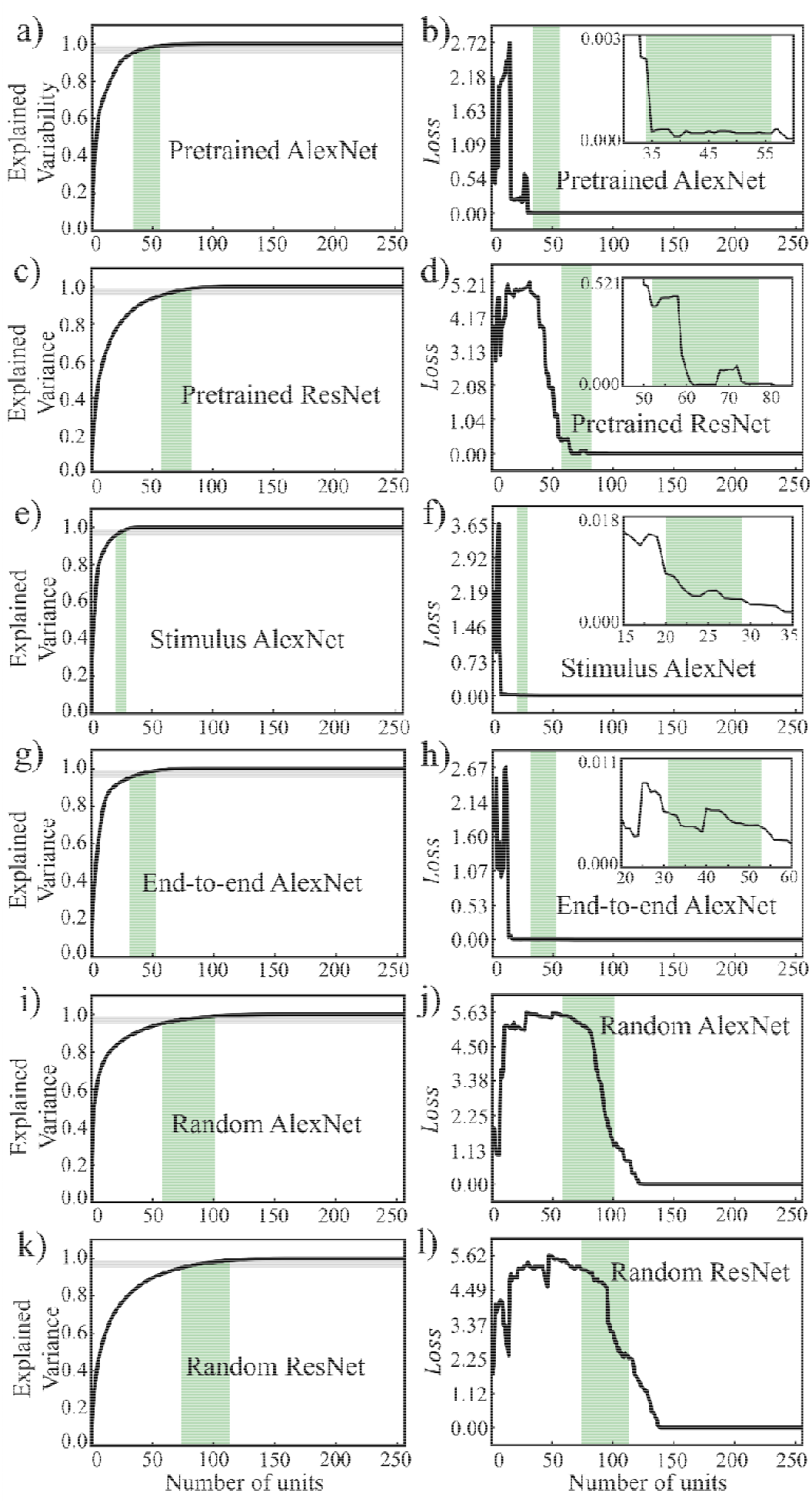
Cumulative variance explained and lesion robustness across network variants. (a,c,e,g,i,k) Cumulative variance in unit activity as a function of the number of recurrent units for each model. The grey horizontal lines marked 95% and 99% explained variance; the green shaded band indicated the corresponding range of unit counts. (b,d,f,h,j,l) Model performance () after virtual lesions as a function of the number of retained units in the advanced cognitive module for the same models. Insets zoomed in on the range spanning the 95–99% bands.

**Fig. S11.**
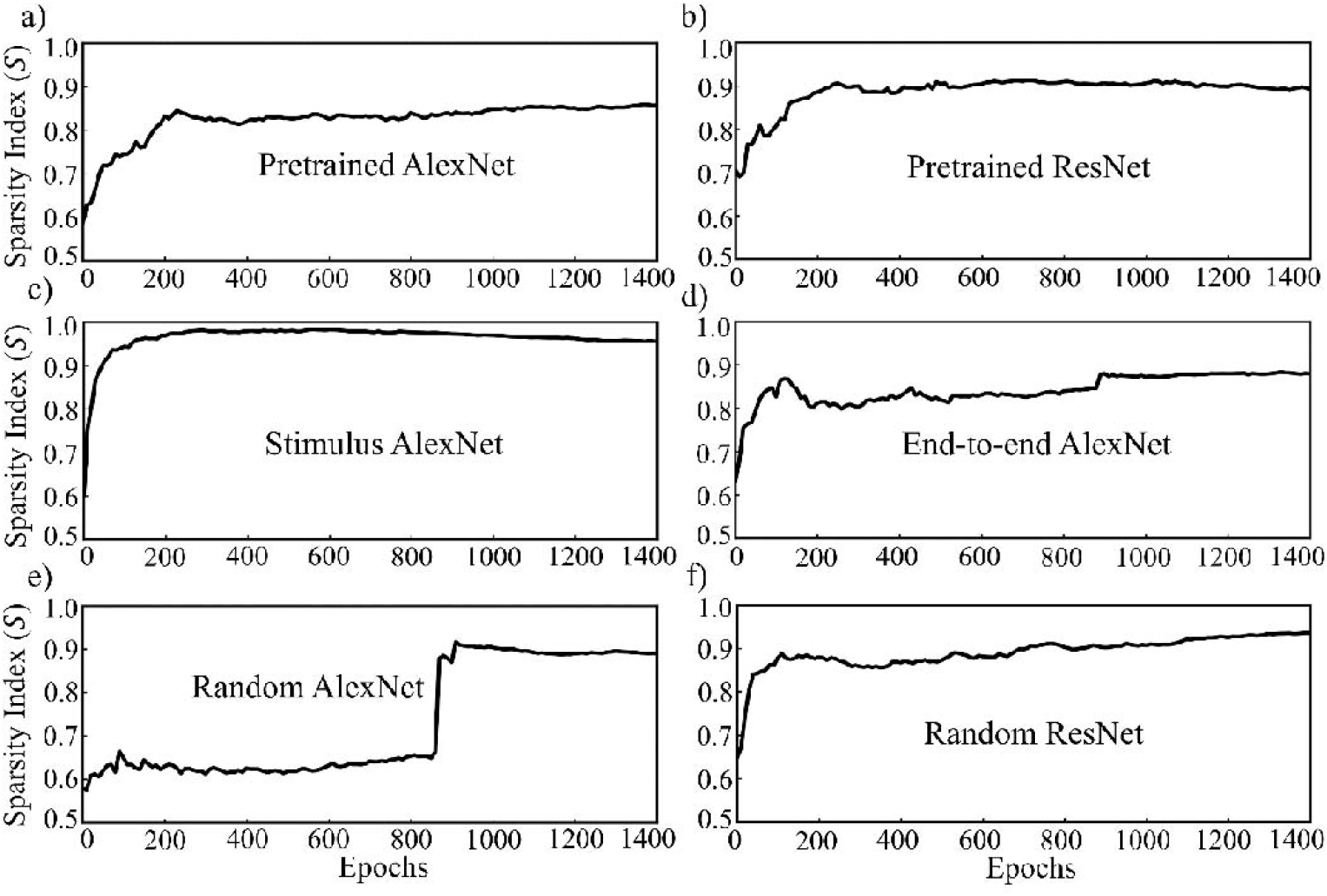
Evolution of the sparsity index of the advanced cognitive module across training epochs. (a) Pretrained AlexNet; (b) Pretrained ResNet; (c) Stimulus AlexNet; (d) End-to-end AlexNet; (e) Random AlexNet; (f) Random ResNet.

**Fig. S12.**
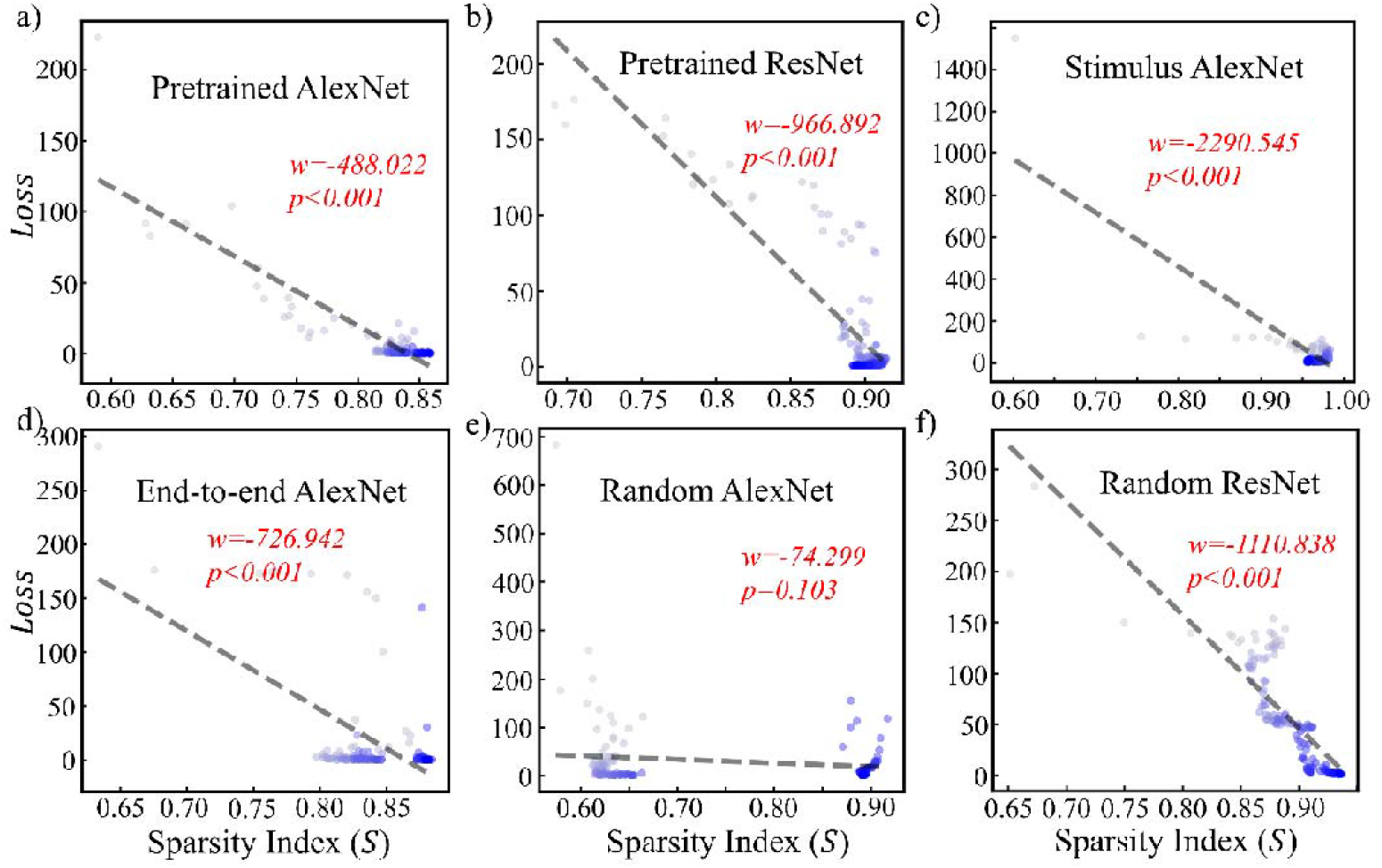
Relationship between model performance () and the sparsity index of the advanced cognitive module. Each point denoted one training epoch, with darker symbols indicating later epochs. Dashed lines showed linear regression fits with slope and corresponding-values.

**Fig. S13.**
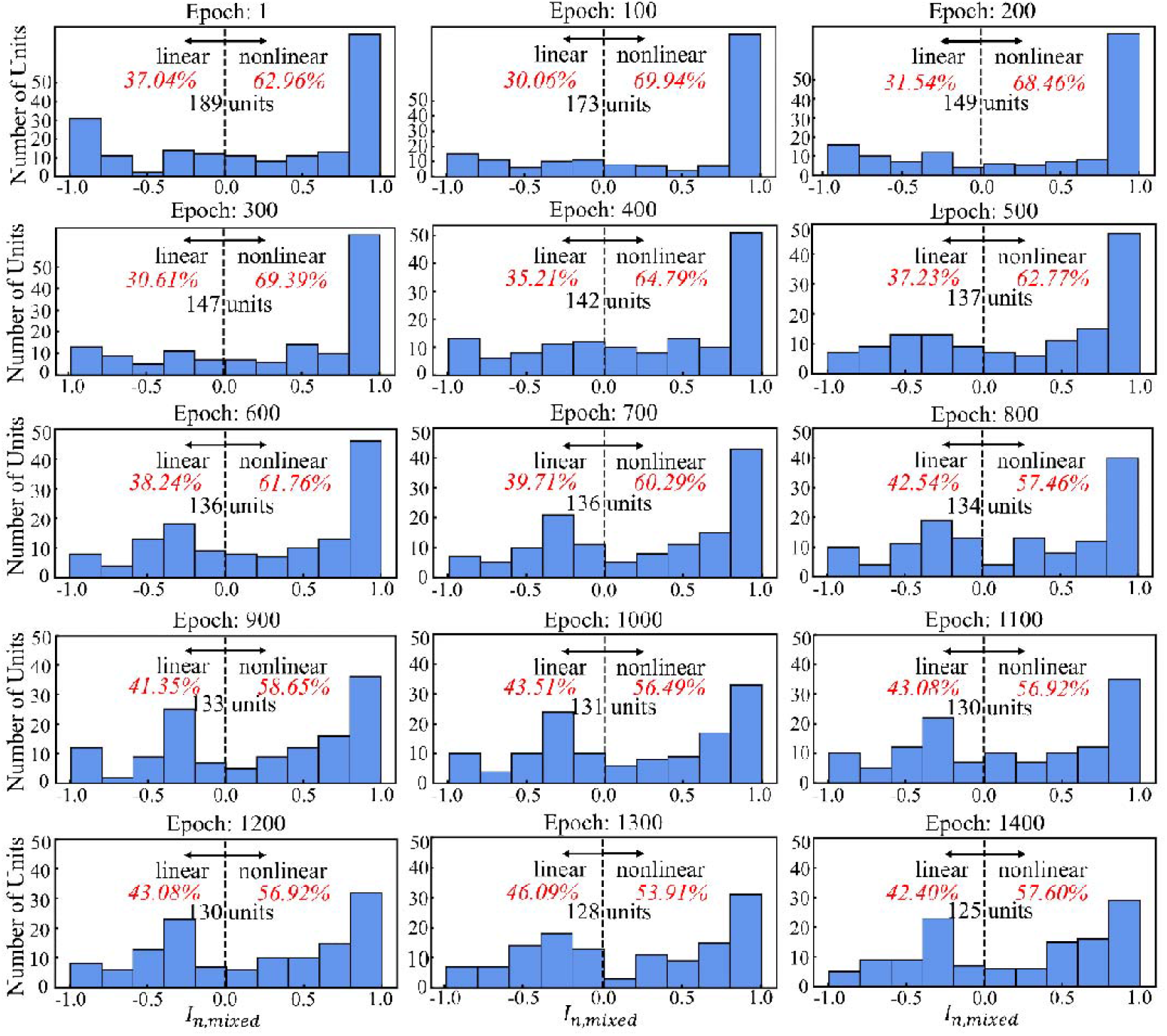
Distribution of unit-wise mixed-selectivity preference () in the advanced cognitive module of the Pretrained AlexNet across training epochs. Red text indicated the proportion of units dominated by linear ( < 0) versus nonlinear ( > 0) tuning, and the total number of mixed-selective units was shown in each panel.

## References and notes

Amengual, J. L., & Ben Hamed, S. (2021). Revisiting Persistent Neuronal Activity During Covert Spatial Attention. Frontiers in Neural Circuits, 15, 679796. 10.3389/fncir.2021.679796

Aoi, M. C., Mante, V., & Pillow, J. W. (2020). Prefrontal cortex exhibits multidimensional dynamic encoding during decision-making. Nature Neuroscience, 23(11), 1410–1420. 10.1038/s41593-020-0696-5

Asplund, C. L., Todd, J. J., Snyder, A. P., & Marois, R. (2010). A central role for the lateral prefrontal cortex in goal-directed and stimulus-driven attention. Nature Neuroscience, 13(4), 507–512. 10.1038/nn.2509

Beyeler, M., Rounds, E. L., Carlson, K. D., Dutt, N., & Krichmar, J. L. (2019). Neural correlates of sparse coding and dimensionality reduction. PLOS Computational Biology, 15(6), e1006908. 10.1371/journal.pcbi.1006908

Chang, L. J., Gianaros, P. J., Manuck, S. B., Krishnan, A., & Wager, T. D. (2015). A Sensitive and Specific Neural Signature for Picture-Induced Negative Affect. PLOS Biology, 13(6), e1002180. 10.1371/journal.pbio.1002180

Cheal, M., & Lyon, D. R. (1991). Central and Peripheral Precuing of Forced-Choice Discrimination. The Quarterly Journal of Experimental Psychology Section A, 43(4), 859–880. 10.1080/14640749108400960

Chung, S., & Abbott, L. F. (2021). Neural population geometry: An approach for understanding biological and artificial neural networks. Current Opinion in Neurobiology, 70, 137–144. 10.1016/j.conb.2021.10.010

Corbetta, M., & Shulman, G. L. (2002). Control of goal-directed and stimulus-driven attention in the brain. Nature Reviews Neuroscience, 3(3), 201–215. 10.1038/nrn755

Di Bello, F., Ceccarelli, F., Messinger, A., & Genovesio, A. (2025). Endogenous and exogenous attentional interplay through mixed prefrontal cortex resources. Current Biology, 35(16), 3825–3838.e3. 10.1016/j.cub.2025.06.070

Driver, J., Davis, G., Ricciardelli, P., Kidd, P., Maxwell, E., & Baron-Cohen, S. (1999). Gaze Perception Triggers Reflexive Visuospatial Orienting. Visual Cognition, 6(5), 509–540. 10.1080/135062899394920

Dubreuil, A., Valente, A., Beiran, M., Mastrogiuseppe, F., & Ostojic, S. (2022). The role of population structure in computations through neural dynamics. Nature Neuroscience, 25(6), 783–794. 10.1038/s41593-022-01088-4

Ebitz, R. B., & Hayden, B. Y. (2021). The population doctrine in cognitive neuroscience. Neuron, 109(19), 3055–3068. 10.1016/j.neuron.2021.07.011

Eichenbaum, H. (2018). Barlow versus Hebb: When is it time to abandon the notion of feature detectors and adopt the cell assembly as the unit of cognition? Neuroscience Letters, 680, 88–93. 10.1016/j.neulet.2017.04.006

Engell, A. D., Nummenmaa, L., Oosterhof, N. N., Henson, R. N., Haxby, J. V., & Calder, A. J. (2010). Differential activation of frontoparietal attention networks by social and symbolic spatial cues. Social Cognitive and Affective Neuroscience, 5(4), 432–440. 10.1093/scan/nsq008

Finn, E. S., Shen, X., Scheinost, D., Rosenberg, M. D., Huang, J., Chun, M. M., Papademetris, X., & Constable, R. T. (2015). Functional connectome fingerprinting: Identifying individuals using patterns of brain connectivity. Nature Neuroscience, 18(11), 1664–1671. 10.1038/nn.4135

Fox, M. D., Corbetta, M., Snyder, A. Z., Vincent, J. L., & Raichle, M. E. (2006). Spontaneous neuronal activity distinguishes human dorsal and ventral attention systems. Proceedings of the National Academy of Sciences, 103(26), 10046–10051. 10.1073/pnas.0604187103

Friesen, C. K., & Kingstone, A. (1998). The eyes have it! Reflexive orienting is triggered by nonpredictive gaze. Psychonomic Bulletin & Review, 5(3), 490–495. 10.3758/BF03208827

Friesen, C. K., & Kingstone, A. (2003). Abrupt onsets and gaze direction cues trigger independent reflexive attentional effects. Cognition, 87(1), B1–B10. 10.1016/S0010-0277(02)00181-6

Frischen, A., & Tipper, S. P. (2004). Orienting attention via observed gaze shift evokes longer term inhibitory effects: Implications for social interactions, attention, and memory. Journal of Experimental Psychology: General, 133(4), 516.

Fusi, S., Miller, E. K., & Rigotti, M. (2016). Why neurons mix: High dimensionality for higher cognition. Current Opinion in Neurobiology, 37, 66–74. 10.1016/j.conb.2016.01.010

Gabay, S., & Behrmann, M. (2014). Attentional dynamics mediated by subcortical mechanisms. Attention, Perception, & Psychophysics, 76(8), 2375–2388. 10.3758/s13414-014-0725-0

Gao, W., Zhu, C., Si, B., Zhou, L., & Zhou, K. (2025). Precision-dependent modulation of social attention. NeuroImage, 310, 121166. 10.1016/j.neuroimage.2025.121166

Gottlieb, J. P., Kusunoki, M., & Goldberg, M. E. (1998). The representation of visual salience in monkey parietal cortex. Nature, 391(6666), 481–484. 10.1038/35135

Graham, D. J., & Field, D. J. (2007). Sparse Coding in the Neocortex. In Evolution of Nervous Systems (Vol. 3, pp. 181–187). Elsevier. 10.1016/B0-12-370878-8/00064-1

Guedj, C., & Vuilleumier, P. (2023). Modulation of pulvinar connectivity with cortical areas in the control of selective visual attention. NeuroImage, 266, 119832. 10.1016/j.neuroimage.2022.119832

Hadjidimitrakis, K., Bakola, S., Wong, Y. T., & Hagan, M. A. (2019). Mixed Spatial and Movement Representations in the Primate Posterior Parietal Cortex. Frontiers in Neural Circuits, 13, 15. 10.3389/fncir.2019.00015

He, K., Zhang, X., Ren, S., & Sun, J. (2016). Deep Residual Learning for Image Recognition. 2016 IEEE Conference on Computer Vision and Pattern Recognition (CVPR), 770–778. 10.1109/CVPR.2016.90

Hietanen, J. K., Nummenmaa, L., Nyman, M. J., Parkkola, R., & Hämäläinen, H. (2006). Automatic attention orienting by social and symbolic cues activates different neural networks: An fMRI study. NeuroImage, 33(1), 406–413. 10.1016/j.neuroimage.2006.06.048

Hodsoll, J., Mevorach, C., & Humphreys, G. W. (2009). Driven to Less Distraction: rTMS of the Right Parietal Cortex Reduces Attentional Capture in Visual Search. Cerebral Cortex, 19(1), 106–114. 10.1093/cercor/bhn070

Hooker, C. I., Paller, K. A., Gitelman, D. R., Parrish, T. B., Mesulam, M.-M., & Reber, P. J. (2003). Brain networks for analyzing eye gaze. Cognitive Brain Research, 17(2), 406–418. 10.1016/S0926-6410(03)00143-5

Jääskeläinen, I. P., Glerean, E., Klucharev, V., Shestakova, A., & Ahveninen, J. (2022). Do sparse brain activity patterns underlie human cognition? NeuroImage, 263, 119633. 10.1016/j.neuroimage.2022.119633

Jingling, L., Lin, H.-F., Tsai, C.-J., & Lin, C.-C. (2015). Development of inhibition of return for eye gaze in adolescents. Journal of Experimental Child Psychology, 137, 76–84. 10.1016/j.jecp.2015.04.001

Joseph, R. M., Fricker, Z., & Keehn, B. (2015). Activation of frontoparietal attention networks by non-predictive gaze and arrow cues. Social Cognitive and Affective Neuroscience, 10(2), 294–301. 10.1093/scan/nsu054

Kadohisa, M., Rolls, E. T., & Verhagen, J. V. (2004). Orbitofrontal cortex: Neuronal representation of oral temperature and capsaicin in addition to taste and texture. Neuroscience, 127(1), 207–221. 10.1016/j.neuroscience.2004.04.037

Kingma, D. P., & Ba, J. (2017). Adam: A Method for Stochastic Optimization (No. arXiv:1412.6980). arXiv. http://arxiv.org/abs/1412.6980

Kingstone, A., Friesen, C. K., & Gazzaniga, M. S. (2000). Reflexive Joint Attention Depends on Lateralized Cortical Connections. Psychological Science, 11(2), 159–166. 10.1111/1467-9280.00232

Kingstone, A., Tipper, C., Ristic, J., & Ngan, E. (2004). The eyes have it!: An fMRI investigation. Brain and Cognition, 55(2), 269–271. 10.1016/j.bandc.2004.02.037

Klein, R. M. (2000). Inhibition of return. Trends in Cognitive Sciences, 4(4), 138–147.

Kreiman, G., Hung, C. P., Kraskov, A., Quiroga, R. Q., Poggio, T., & DiCarlo, J. J. (2006). Object Selectivity of Local Field Potentials and Spikes in the Macaque Inferior Temporal Cortex. Neuron, 49(3), 433–445. 10.1016/j.neuron.2005.12.019

Krishnan, A., Woo, C.-W., Chang, L. J., Ruzic, L., Gu, X., López-Solà, M., Jackson, P. L., Pujol, J., Fan, J., & Wager, T. D. (2016). Somatic and vicarious pain are represented by dissociable multivariate brain patterns. eLife, 5, e15166. 10.7554/eLife.15166

Krizhevsky, A., Sutskever, I., & Hinton, G. E. (2017). ImageNet classification with deep convolutional neural networks. Communications of the ACM, 60(6), 84–90. 10.1145/3065386

Langdon, C., Genkin, M., & Engel, T. A. (2023). A unifying perspective on neural manifolds and circuits for cognition. Nature Reviews Neuroscience, 24(6), 363–377. 10.1038/s41583-023-00693-x

Langton, S. R. H., & Bruce, V. (1999). Reflexive Visual Orienting in Response to the Social Attention of Others. Visual Cognition, 6(5), 541–567. 10.1080/135062899394939

Lockhofen, D. E. L., Gruppe, H., Ruprecht, C., Gallhofer, B., & Sammer, G. (2014). Hemodynamic Response Pattern of Spatial Cueing is Different for Social and Symbolic Cues. Frontiers in Human Neuroscience, 8. 10.3389/fnhum.2014.00912

Lundqvist, M., Brincat, S. L., Rose, J., Warden, M. R., Buschman, T. J., Miller, E. K., & Herman, P. (2023). Working memory control dynamics follow principles of spatial computing. Nature Communications, 14(1), 1429. 10.1038/s41467-023-36555-4

Masse, N. Y., Yang, G. R., Song, H. F., Wang, X.-J., & Freedman, D. J. (2019). Circuit mechanisms for the maintenance and manipulation of information in working memory. Nature Neuroscience, 22(7), 1159–1167. 10.1038/s41593-019-0414-3

McKenzie, S., Keene, C. S., Farovik, A., Bladon, J., Place, R., Komorowski, R., & Eichenbaum, H. (2016). Representation of memories in the cortical–hippocampal system: Results from the application of population similarity analyses. Neurobiology of Learning and Memory, 134, 178–191. 10.1016/j.nlm.2015.12.008

Meyer, K. N., Du, F., Parks, E., & Hopfinger, J. B. (2018). Exogenous vs. endogenous attention: Shifting the balance of fronto-parietal activity. Neuropsychologia, 111, 307–316. 10.1016/j.neuropsychologia.2018.02.006

Młynarski, W., & Tkačik, G. (2022). Efficient coding theory of dynamic attentional modulation. PLOS Biology, 20(12), e3001889. 10.1371/journal.pbio.3001889

Mulckhuyse, M., & Theeuwes, J. (2010). Unconscious attentional orienting to exogenous cues: A review of the literature. Acta Psychologica, 134(3), 299–309. 10.1016/j.actpsy.2010.03.002

Muller, H. J., & Rabbitt, P. M. A. (1989). Reflexive and Voluntary Orienting of Visual Attention: Time Course of Activation and Resistance to Interruption. J Exp Psychol Hum Percept Perform, 15(2), 315–330. 10.1037//0096-1523.15.2.315

Pagan, M., Tang, V. D., Aoi, M. C., Pillow, J. W., Mante, V., Sussillo, D., & Brody, C. D. (2025). Individual variability of neural computations underlying flexible decisions. Nature, 639(8054), 421–429. 10.1038/s41586-024-08433-6

Palm, G. (2013). Neural associative memories and sparse coding. Neural Networks, 37, 165–171. 10.1016/j.neunet.2012.08.013

Panzeri, S., Macke, J. H., Gross, J., & Kayser, C. (2015). Neural population coding: Combining insights from microscopic and mass signals. Trends in Cognitive Sciences, 19(3), 162–172. 10.1016/j.tics.2015.01.002

Parthasarathy, A., Herikstad, R., Bong, J. H., Medina, F. S., Libedinsky, C., & Yen, S.-C. (2017). Mixed selectivity morphs population codes in prefrontal cortex. Nature Neuroscience, 20(12), 1770–1779. 10.1038/s41593-017-0003-2

Peelen, M. V., Heslenfeld, D. J., & Theeuwes, J. (2004). Endogenous and exogenous attention shifts are mediated by the same large-scale neural network. NeuroImage, 22(2), 822–830. 10.1016/j.neuroimage.2004.01.044

Perrodin, C., Kayser, C., Logothetis, N. K., & Petkov, C. I. (2011). Voice Cells in the Primate Temporal Lobe. Current Biology, 21(16), 1408–1415. 10.1016/j.cub.2011.07.028

Petersen, S. E., & Posner, M. I. (2012). The Attention System of the Human Brain: 20 Years After. Annual Review of Neuroscience, 35(1), 73–89. 10.1146/annurev-neuro-062111-150525

Piwek, E. P., Stokes, M. G., & Summerfield, C. (2023). A recurrent neural network model of prefrontal brain activity during a working memory task. PLOS Computational Biology, 19(10), e1011555. 10.1371/journal.pcbi.1011555

Posner, M. I. (1980). Orienting of Attention. Quarterly Journal of Experimental Psychology, 32(1), 3–25. 10.1080/00335558008248231

Posner, M. I., & Cohen, Y. (1984). Components of visual orienting. Attention and Performance X: Control of Language Processes, 32, 531–556.

Raposo, D., Kaufman, M. T., & Churchland, A. K. (2014). A category-free neural population supports evolving demands during decision-making. Nature Neuroscience, 17(12), 1784–1792. 10.1038/nn.3865

Reinert, S., Hübener, M., Bonhoeffer, T., & Goltstein, P. M. (2021). Mouse prefrontal cortex represents learned rules for categorization. Nature, 593(7859), 411–417. 10.1038/s41586-021-03452-z

Richards, B. A., Lillicrap, T. P., Beaudoin, P., Bengio, Y., Bogacz, R., Christensen, A., Clopath, C., Costa, R. P., de Berker, A., Ganguli, S., Gillon, C. J., Hafner, D., Kepecs, A., Kriegeskorte, N., Latham, P., Lindsay, G. W., Miller, K. D., Naud, R., Pack, C. C., … Kording, K. P. (2019). A deep learning framework for neuroscience. Nature Neuroscience, 22(11), 1761–1770. 10.1038/s41593-019-0520-2

Riggio, L., & Kirsner, K. (1997). The relationship between central cues and peripheral cues in covert visual orientation. Perception & Psychophysics, 59(6), 885–899. 10.3758/BF03205506

Rigotti, M., Barak, O., Warden, M. R., Wang, X.-J., Daw, N. D., Miller, E. K., & Fusi, S. (2013). The importance of mixed selectivity in complex cognitive tasks. Nature, 497(7451), 585–590. 10.1038/nature12160

Roseboom, W., Fountas, Z., Nikiforou, K., Bhowmik, D., Shanahan, M., & Seth, A. K. (2019). Activity in perceptual classification networks as a basis for human subjective time perception. Nature Communications, 10(1), 267. 10.1038/s41467-018-08194-7

Rosenberg, M. D., Finn, E. S., Scheinost, D., Constable, R. T., & Chun, M. M. (2017). Characterizing Attention with Predictive Network Models. Trends in Cognitive Sciences, 21(4), 290–302. 10.1016/j.tics.2017.01.011

Rosenberg, M. D., Finn, E. S., Scheinost, D., Papademetris, X., Shen, X., Constable, R. T., & Chun, M. M. (2016). A neuromarker of sustained attention from whole-brain functional connectivity. Nature Neuroscience, 19(1), 165–171. 10.1038/nn.4179

Rossi, A. F., Bichot, N. P., Desimone, R., & Ungerleider, L. G. (2007). Top Down Attentional Deficits in Macaques with Lesions of Lateral Prefrontal Cortex. Journal of Neuroscience, 27(42), 11306–11314. 10.1523/JNEUROSCI.2939-07.2007

Saez, A., Rigotti, M., Ostojic, S., Fusi, S., & Salzman, C. D. (2015). Abstract Context Representations in Primate Amygdala and Prefrontal Cortex. Neuron, 87(4), 869–881. 10.1016/j.neuron.2015.07.024

Samuel, A. G., & Kat, D. (2003). Inhibition of return: A graphical meta-analysis of its time course and an empirical test of its temporal and spatial properties. Psychonomic Bulletin & Review, 10(4), 897–906. 10.3758/BF03196550

Sapountzis, P., Paneri, S., Papadopoulos, S., & Gregoriou, G. G. (2022). Dynamic and stable population coding of attentional instructions coexist in the prefrontal cortex. Proceedings of the National Academy of Sciences, 119(40), e2202564119. 10.1073/pnas.2202564119

Sato, W., Kochiyama, T., Uono, S., & Toichi, M. (2016). Neural mechanisms underlying conscious and unconscious attentional shifts triggered by eye gaze. NeuroImage, 124, 118–126. 10.1016/j.neuroimage.2015.08.061

Sato, W., Kochiyama, T., Uono, S., & Yoshikawa, S. (2009). Commonalities in the neural mechanisms underlying automatic attentional shifts by gaze, gestures, and symbols. NeuroImage, 45(3), 984–992. 10.1016/j.neuroimage.2008.12.052

Saxena, S., & Cunningham, J. P. (2019). Towards the neural population doctrine. Current Opinion in Neurobiology, 55, 103–111. 10.1016/j.conb.2019.02.002

Sheahan, H., Luyckx, F., Nelli, S., Teupe, C., & Summerfield, C. (2021). Neural state space alignment for magnitude generalization in humans and recurrent networks. Neuron, 109(7), 1214–1226.e8. 10.1016/j.neuron.2021.02.004

Shi, B., Song, Y., Joshi, N., Darrell, T., & Wang, X. (2022, June 12). Visual Attention Emerges from Recurrent Sparse Reconstruction. arXiv. International Conference on Machine Learning.

Shulman, G. L., Pope, D. L. W., Astafiev, S. V., McAvoy, M. P., Snyder, A. Z., & Corbetta, M. (2010). Right Hemisphere Dominance during Spatial Selective Attention and Target Detection Occurs Outside the Dorsal Frontoparietal Network. Journal of Neuroscience, 30(10), 3640–3651. 10.1523/JNEUROSCI.4085-09.2010

Song, K., Meng, M., Chen, L., Zhou, K., & Luo, H. (2014). Behavioral Oscillations in Attention: Rhythmic α Pulses Mediated through θ Band. The Journal of Neuroscience, 34(14), 4837–4844. 10.1523/JNEUROSCI.4856-13.2014

Stokes, M. G., Kusunoki, M., Sigala, N., Nili, H., Gaffan, D., & Duncan, J. (2013). Dynamic Coding for Cognitive Control in Prefrontal Cortex. Neuron, 78(2), 364–375. 10.1016/j.neuron.2013.01.039

Tavor, I., Jones, O. P., Mars, R. B., Smith, S. M., Behrens, T. E., & Jbabdi, S. (2016). Task-free MRI predicts individual differences in brain activity during task performance. Science, 352(6282), 216–220. 10.1126/science.aad8127

Tipper, C. M., Handy, T. C., Giesbrecht, B., & Kingstone, A. (2008). Brain Responses to Biological Relevance. Journal of Cognitive Neuroscience, 20(5), 879–891. 10.1162/jocn.2008.20510

Uono, S., Sato, W., & Kochiyama, T. (2014). Commonalities and differences in the spatiotemporal neural dynamics associated with automatic attentional shifts induced by gaze and arrows. Neuroscience Research, 87, 56–65. 10.1016/j.neures.2014.07.003

Vecera, S. P., & Rizzo, M. (2004). What are you looking at? Neuropsychologia, 42(12), 1657–1665. 10.1016/j.neuropsychologia.2004.04.009

Vecera, S. P., & Rizzo, M. (2006). Eye gaze does not produce reflexive shifts of attention: Evidence from frontal-lobe damage. Neuropsychologia, 44(1), 150–159. 10.1016/j.neuropsychologia.2005.04.010

Vinje, W. E., & Gallant, J. L. (2000). Sparse Coding and Decorrelation in Primary Visual Cortex During Natural Vision. Science, 287(5456), 1273–1276. 10.1126/science.287.5456.1273

Vossel, S., Geng, J. J., & Fink, G. R. (2014). Dorsal and Ventral Attention Systems: Distinct Neural Circuits but Collaborative Roles. The Neuroscientist, 20(2), 150–159. 10.1177/1073858413494269

Vossel, S., Mathys, C., Daunizeau, J., Bauer, M., Driver, J., Friston, K. J., & Stephan, K. E. (2014). Spatial Attention, Precision, and Bayesian Inference: A Study of Saccadic Response Speed. Cerebral Cortex, 24(6), 1436–1450. 10.1093/cercor/bhs418

Vyas, S., Golub, M. D., Sussillo, D., & Shenoy, K. V. (2020). Computation Through Neural Population Dynamics. Annual Review of Neuroscience, 43(1), 249–275. 10.1146/annurev-neuro-092619-094115

Wang, R., Yuan, T., Wang, L., & Jiang, Y. (2024). A common and specialized neural code for social attention triggered by eye gaze and biological motion. NeuroImage, 301, 120889. 10.1016/j.neuroimage.2024.120889

Wixted, J. T., Squire, L. R., Jang, Y., Papesh, M. H., Goldinger, S. D., Kuhn, J. R., Smith, K. A., Treiman, D. M., & Steinmetz, P. N. (2014). Sparse and distributed coding of episodic memory in neurons of the human hippocampus. Proceedings of the National Academy of Sciences, 111(26), 9621–9626. 10.1073/pnas.1408365111

Wu, E. X. W., Liaw, G. J., Goh, R. Z., Chia, T. T. Y., Chee, A. M. J., Obana, T., Rosenberg, M. D., Yeo, B. T. T., & Asplund, C. L. (2020). Overlapping attentional networks yield divergent behavioral predictions across tasks: Neuromarkers for diffuse and focused attention? NeuroImage, 209, 116535. 10.1016/j.neuroimage.2020.116535

Xu, S., Zhang, S., & Geng, H. (2011). Gaze-induced joint attention persists under high perceptual load and does not depend on awareness. Vision Research, 51(18), 2048–2056. 10.1016/j.visres.2011.07.023

Yang, A., Tian, J., Wang, W., Liu, J., Zhou, L., & Zhou, K. (2025). Shared and distinct neural signatures of feature and spatial attention. NeuroImage. 10.1016/j.neuroimage.2025.121296

Yang, G. R., Joglekar, M. R., Song, H. F., Newsome, W. T., & Wang, X.-J. (2019). Task representations in neural networks trained to perform many cognitive tasks. Nature Neuroscience, 22(2), 297–306.

Ye, Z., Li, H., Tian, L., & Zhou, C. (2025). The effects of the post-delay epochs on working memory error reduction. PLOS Computational Biology, 21(5), e1013083. 10.1371/journal.pcbi.1013083

Yuste, R. (2015). From the neuron doctrine to neural networks. Nature Reviews Neuroscience, 16(8), 487–497. 10.1038/nrn3962

Zhang, C. Y., Aflalo, T., Revechkis, B., Rosario, E., Ouellette, D., Pouratian, N., & Andersen, R. A. (2020). Preservation of Partially Mixed Selectivity in Human Posterior Parietal Cortex across Changes in Task Context. Eneuro, 7(2), ENEURO.0222-19.2019. 10.1523/ENEURO.0222-19.2019

Zhang, C. Y., Aflalo, T., Revechkis, B., Rosario, E. R., Ouellette, D., Pouratian, N., & Andersen, R. A. (2017). Partially Mixed Selectivity in Human Posterior Parietal Association Cortex. Neuron, 95(3), 697–708.e4. 10.1016/j.neuron.2017.06.040

Zhou, F., Zhao, W., Qi, Z., Geng, Y., Yao, S., Kendrick, K. M., Wager, T. D., & Becker, B. (2021). A distributed fMRI-based signature for the subjective experience of fear. Nature Communications, 12(1), 6643. 10.1038/s41467-021-26977-3

